# Optical Modulation of Blood-Brain-Tumor Barrier Permeability Enhances Drug Delivery in Diverse Preclinical Glioblastoma Models

**DOI:** 10.1101/2022.06.07.495027

**Authors:** Qi Cai, Xiaoqing Li, Hejian Xiong, Hanwen Fan, Xiaofei Gao, Vamsidhara Vemireddy, Ryan Margolis, Junjie Li, Xiaoqian Ge, Monica Giannotta, Elisabetta Dejana, Kenneth Hoyt, Robert Bachoo, Zhenpeng Qin

**Affiliations:** Department of Mechanical Engineering, University of Texas at Dallas, Richardson, TX 75080, USA; Department of Bioengineering, University of Texas at Dallas, Richardson, TX 75080, USA; Department of Internal Medicine, University of Texas Southwestern Medical Center, Dallas, TX 75390, USA; Harold C. Simmons Comprehensive Cancer Center, University of Texas Southwestern Medical Center, Dallas, TX 75390, USA; FIRC Institute of Molecular Oncology Foundation (IFOM), 20139 Milan, Italy; Department of Neurology, University of Texas Southwestern Medical Center, Dallas, TX 75390, USA; Department of Surgery, University of Texas Southwestern Medical Center, Dallas, TX 75390, USA; Center for Advanced Pain Studies, University of Texas at Dallas, Richardson, TX 75080, USA

## Abstract

Glioblastoma multiforme (GBM) is the most prevalent malignant tumor in the central nervous system. It has diverse phenotypes, including diffuse single-cell infiltration in which the tumor cells co-opt the normal microvasculature, and the neovascularization of an expanding tumor mass. The blood-brain-tumor barrier (BBTB) is a significant obstacle to GBM treatment and restricts entry of most FDA-approved effective oncology drugs. Herein, we report that picosecond laser excitation of vascular-targeted plasmonic gold nanoparticles (AuNPs) can non-invasively and reversibly modulate the BBTB permeability (optoBBTB). OptoBBTB enhances the delivery of paclitaxel (Taxol) in two genetically engineered glioma models (GEMM) that span the spectrum of GBM phenotypes. OptoBBTB followed by Taxol delivery effectively suppresses tumor growth and prolongs the survival time of both GEMM. Moreover, our results raise the possibility that paclitaxel, which is amongst the most widely used oncology drugs because of its proven efficacy but has been abandoned for GBM following its failure to efficacy in early phase clinical trials due to poor blood-brain barrier (BBB) penetration, could now be reconsidered in combination with strategies to increase BBB permeability. In summary, optoBBTB is a novel and effective approach to increase the delivery of therapeutics with limited BBB permeability to treat neoplastic and non-neoplastic brain diseases.

## Introduction

World Health Organization (WHO) grade IV astrocytoma (Isocitrate Dehydrogenase, IDH, wild-type), known as Glioblastoma multiforme (GBM), is one of the most malignant and aggressive types of primary brain tumor (*1–3*). Despite an aggressive standard of care regimen, including maximal safe resection of the contrast-enhancing regions of T1-weighted MR images (MRI), adjuvant fractionated radiation and concurrent temozolomide (TMZ) chemotherapy, the median overall survival for GBM patients remains abysmal at 12-18 months. GBM recurrence most often (70-80% of cases) occurs in the vicinity, within 1-3cm of the surgical margin. This region is well within the radiation field and in the absence of gadolinium enhancement and is presumed to represent a region of intact BBB. Therefore, one of the challenges in GBM treatment is the delivery of chemotherapeutics at effective levels to residual cells at the margins of the surgical resection that are protected by an intact BBB (*4*). Formed by a tight-junction (TJ) complex and adherens junctions and modulated by surrounding stromal cells such as pericytes and astrocytes, the BBB excludes the delivery of 98% of small-molecule and nearly all large-molecule drugs to subtherapeutic levels (*5*). Driven by high levels of angiogenic signals, GBM cells are capable of inducing microvascular proliferation which results in irregular, ectatic structures, including glomeruloid and end-vessels, with limited BBB function. Within the same tumor, GBM cells at the tumor margins are characterized by diffuse single-cell infiltration through the brain parenchyma including in neuron-rich regions of grey matter neuropil and along white matter tracts (*6, 7*). Here, GBM cells co-opt the pre-existing dense brain microvasculature for metabolic support and nutrient exchange without disrupting the normal structure or functions of the BBB (*8–10*). In view of the spatial heterogeneity of BBB integrity and especially following the complete resection, when the scant remaining tumor cells are restricted to the tumor margin regions of intact BBB, effective drug delivery to the tumor at therapeutic concentrations remains an unmet challenge.

Several approaches to overcome the BBTB for therapeutic delivery have been developed, including but not limited to disrupting the BBB using mannitol, opening the tight junction by tight junction modulator, and enhancing drug penetration through inhibiting drug efflux transporters or via receptor-mediated transport (*11–14*). While these strategies may improve drug delivery to brain tumors, high incidences of complications and potential for toxicity have impeded progress in the clinical translation of these BBB-disrupting technologies (*11, 14, 15*). Focused ultrasound (FUS) in combination with intravenously (i.v.) administered microbubbles is a local, minimally invasive method for transiently disrupting the BBTB disruption, which has progressed to early-phase clinical trials (*16–18*). However, knowledge gaps remain, such as optimal parameters for sonication and choice of the size of the microbubbles. Furthermore, many of the preclinical in vivo studies of intracranial xenograft models generated with long-established cell lines are often poor representations of GBM genotype and phenotype (*19*). Indeed, numerous promising preclinical therapeutic strategies have shown limited efficacy or failed at the clinical trial stage, highlighting the compelling need to exploit and test new more rigorous preclinical models to bridge the gap between preclinical efficacy and successful clinical translation (*20–22*).

Here, we report a non-invasive and reversible BBTB modulation approach (optoBBTB), utilizing two GEMM models that define two major microvascular phenotypes seen in de novo GBM. These phenotypes include regions of diffuse single-cell infiltration where tumor cells co-opt the normal microvasculature for metabolic support, and regions of microvascular proliferation with compromised BBB functions. Our previous study showed that picosecond (ps) laser stimulation of TJ targeted gold nanoparticles (AuNPs) can non-invasively and reversibly increase the BBB permeability to facilitate the delivery of antibodies, adeno-associated virus, and liposomes into the brain parenchyma (*23*). Adopting this versatile platform, we demonstrate a novel approach to modulate the BBTB permeability locally for anti-cancer drug delivery. This study reveals that following repeated cycles of optoBBTB and systemic administration of paclitaxel (Taxol) suppress the tumor growth by reducing tumor cell proliferation and increasing cell death, and result in a significantly improved median survival in both GBM xenograft models. Therefore, the optoBBTB approach offers high efficacy, safety and spatiotemporal precision for brain drug delivery and is promising for further evaluation in the clinical setting following successful large animal studies.

## Results

### Analysis of the characteristics of human GBM

First, we analyzed the human GBM characteristics to serve as guidance for selecting appropriate glioma mouse models. Human GBM is characterized by infiltrative growth, high mitotic activity in the tumor core, and hemorrhagic regions by Hematoxylin and eosin (H&E) staining (**Fig. S1A**). The enrichment of cell nuclei was further confirmed by Hoechst staining (**Fig. S1B**). Immunohistochemical (IHC) staining shows CD31+ blood vessels with disorganized and ectatic structures in the tumor core compared to the tumor margin (**Fig. S1B**), suggesting the *de novo* angiogenesis through the extension of the nearby vessels. To investigate the changes of TJ properties in human GBM sample, we selected zonula occludens 1 (ZO-1) and occludin as two representative proteins, since previous research discovered that TJ components ZO and transmembrane proteins such as occludin, claudins and junctional adhesion molecules (JAM), seal the paracellular route between opposing brain microvascular endothelial cells and regulate the low paracellular permeability of the BBB (*24*). Interestingly, loss of the immunofluorescence of ZO-1 and occludin was observed in the tumor core rather than tumor margin (**Fig. S1B**), indicating a heterogeneous breakdown of functional TJ in human GBM. The above results suggest that human GBM shows a highly vascularized tumor core with disrupted BBTB partially due to the loss of ZO-1 and occludin. In contrast, the infiltrative and intact margin shows strong ZO-1 and occludin signals.

### OptoBBTB improves drug penetration in a diffusely infiltrative GEMM GBM model

As the standard of care, following complete surgical resection of the T1W gadolinium-enhancing regions, residual tumor cells in the surrounding margin regions are protected by an intact BBB (*25, 26*). Our previous work identified a primary conditional astrocyte mouse cell line that carried *lsl.Braf^V600Ef/+^*, *Ink4ab/Arf ^f/f^* and *Pten^f/f^* mutations, when induced to undergo recombination following adeno-Cre infection (*Braf^V600+/-^*, *Ink4ab/Arf^-/,^ Pten^-/-^*, and engineered to express GFP fluorescent, here after referred to as PS5A1, see discussion and supplementary information for details) resulted in a diffusely infiltrative robust glioma model. The BBTB integrity of the mice during PS5A1 glioma progression was first analyzed using i.v. injection of EZ-link biotin (660 Da) and Evans blue (66 kDa, albumin-bound). **Fig. 1A and B** show the dye confinement in the blood vessels in both tumor core and margin at 14, 28, and 42 dpi, and therefore the BBTB remains intact. The PS5A1 tumor cells were seen to display a heterogeneous pattern of brain infiltration, moving along brain capillaries, through the neuropil-rich gray matter, and parallel to myelinated axons along the white matter tracts. Moreover, there was no evidence of angiogenesis with this diffusely invasive glioma model (**Fig. 1C-D**). Vascular density identified by both structure (endothelial marker CD31 positive cells) and perfusion (luminal wall labeling with tomato lectin594) was equivalent in the tumor core and margin regions to that of the contralateral hemisphere (**Fig. 1C-D**). This data suggest that tumor growth and infiltration are promoted by co-opting the normal dense brain microvasculature for nutrient and metabolic support. Moreover, IHC staining showed that JAM-A coverage on blood vessels was consistent in the contralateral side, tumor core, and tumor margin, allowing the anchoring of JAM-A targeted AuNPs (**Fig. 1E**). Taken together, our histopathological characterization of the PS5A1 glioma model suggests that it is a suitable in vivo model for studying the challenges of drug delivery across an intact BBB similar to those encountered at the invasive tumor margin in the clinical setting.

**Fig.1.**
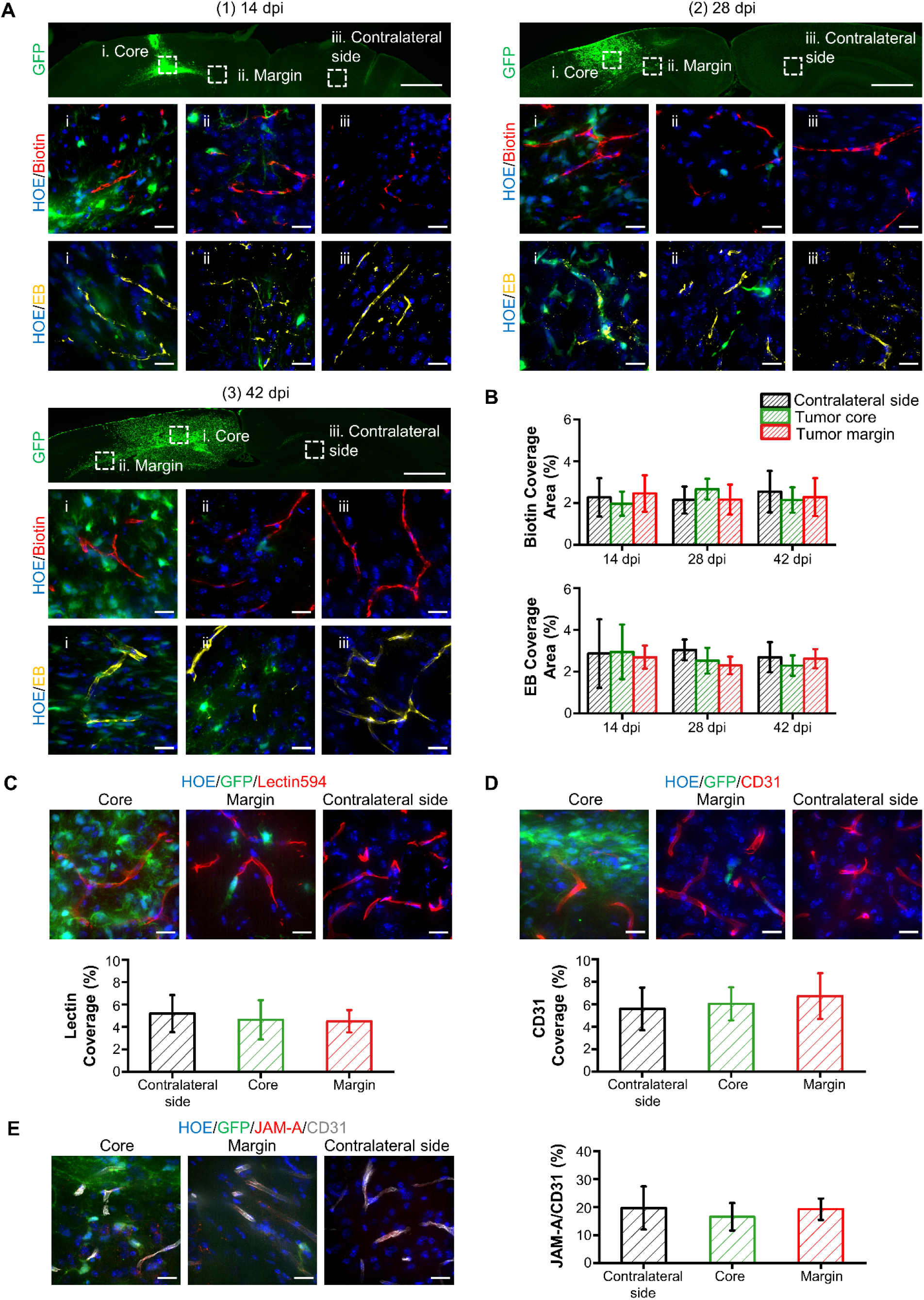
PS5A1 glioma model is infiltrative and has intact BBTB during disease progression. (A) BBTB permeability analysis by using EZ-link biotin (Biotin, red, 660 Da) and Evans blue (EB, yellow, 66 kDa when bound to albumin) at 14, 28, and 42 days post-injection (dpi). The cell nuclei were indicated by Hoechst staining (HOE, blue). The ROIs selected are (i) tumor core, (ii) tumor margin, and (iii) contralateral side with no tumor. The scale bar represents 1 mm in the upper panel and 20 µm in the bottom panels. (B) The quantification of biotin and Evans blue coverage by area fraction. Data was expressed as mean ±SD. One-way Anova, ***p<0.001, N=15 images from 3 mice. (C) Blood vessel labeling with tomato lectin594 and quantification of lectin coverage by area fraction. The scale bar represents 20 µm. One-way Anova, no significant difference. N=10 images. (D) Blood vessel labeling with CD31 and quantification of CD31 coverage by area fraction. The scale bar represents 20 µm. One-way Anova, no significant difference, N=10 images. (E) IHC staining and quantification of JAM-A in PS5A1 GBM model at 14 dpi. The scale bar represents 20 µm. One-way Anova, no significant difference. N=10 images.

Next, we investigated the BBTB modulation using optoBBTB in PS5A1 tumor-bearing mice adopting a two-step process. Firstly, we prepared AuNPs were prepared and i.v. injected into a tumor-bearing mouse to target the TJ component. Next, fluorescent dyes or fluorescence-labeled Taxol was administrated to assess the BBB permeability and the brain uptake, and a transcranial picosecond laser pulse was delivered to the tumor region to stimulate the AuNPs for BBTB modulation (**Fig. 2A**). JAM-A targeted nanoparticles (AuNP-BV11, 50 nm) were selected in this study after characterizing their physicochemical properties (**Fig. S2A-D**). Their ability to modulate BBB and safety profiles have been thoroughly investigated and previously reported (*23*). We then optimized the optoBBTB for the PS5A1 glioma model and tested the therapeutic efficacy. A series of nanoparticle doses and laser fluences were tested (**Table S1)**. 18.5 µg/g of AuNP-BV11 injection followed by 40 mJ/cm^2^ laser fluence were selected for BBTB opening since it showed high opening efficacy with minimized nanoparticle injection (**Fig. S3A)**. We further demonstrated that optoBBTB allowed the delivery of molecules of different sizes, such as EZ-link biotin (660 Da) and Evans blue/albumin (66 kDa) (**Fig. 2B**). The BBTB opening was reversible and gradually recovered in 3 days (**Fig. 3B**).

**Fig. 2.**
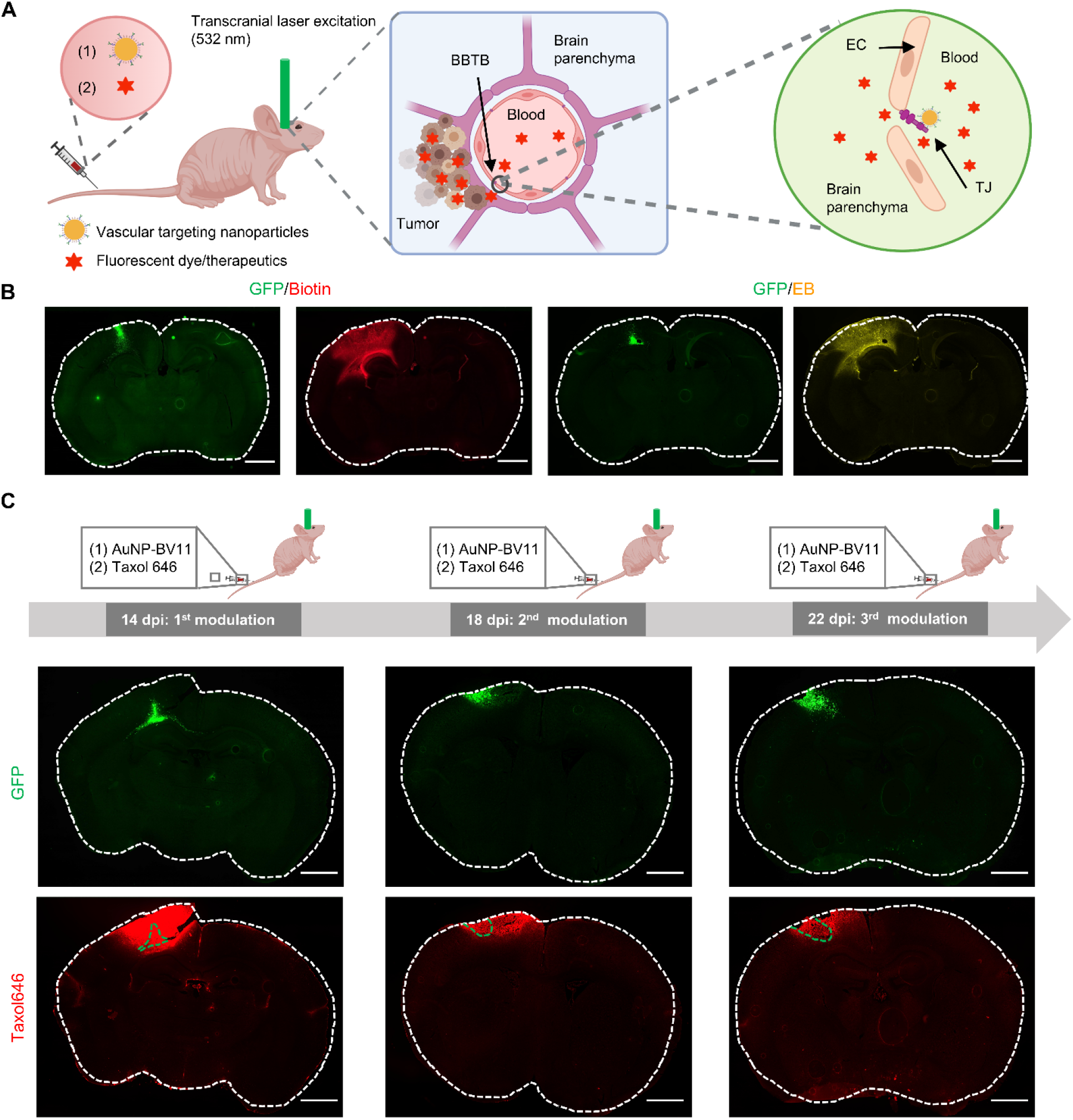
OptoBBTB improves drug penetration in PS5A1 glioma model. (A) Schematic illustration of optoBBTB. (B) Delivery of EZ-link biotin and Evans blue after optoBBTB. The tumor was indicated by GFP fluorescent (GFP), and BBTB opening was indicated by EZ-link biotin (Biotin) or Evans blue leakage (EB). The scale bar represents 1 mm. (C) Multiple BBTB modulations in PS5A1 GBM model at 14, 18, and 22 dpi. AuNP-BV11 and taxol646 were injected intravenously. The tumor cells were indicated by GFP and taxol646 was indicated by red. The scale bar represents 1 mm.

**Fig. 3.**
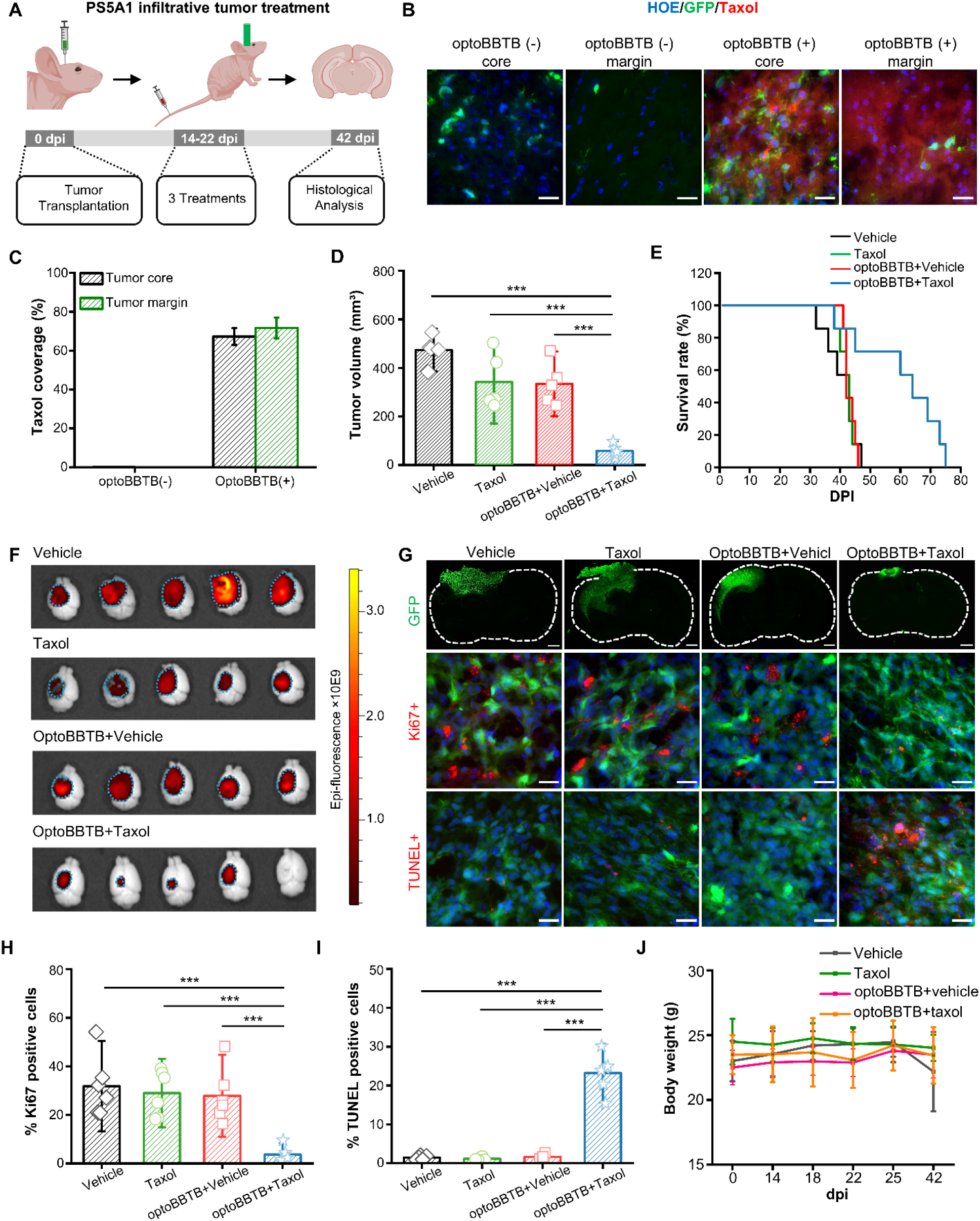
OptoBBTB improves therapeutic outcomes in the infiltrative PS5A1 GBM model. (A) Schematic illustration of the treatment plan. (B) OptoBBTB facilitated the delivery of fluorescent Taxol to tumor core and margin. The scale bar represents 20 µm. (C) Quantification of taxol delivery by fluorescent area fraction. (D) The analysis of tumor volume was measured by GFP fluorescent signal at 42 dpi. One-way Anova, ***p<0.001, N=5 mice in each group. (E) Kaplan-Meier survival analysis, N=7 mice in each group. (F) Fluorescent imaging of tumor size. (G) Top: Tumor size indicated by GFP fluorescent at 42 dpi. The scale bar represents 1mm. Middle: Ki67 staining shows cell proliferation, Bottom: TUNEL staining indicates cell apoptosis. The scale bar represents 20 µm. (H-I) Quantification of Ki67 staining and TUNEL staining after PS5A1 GBM treatment. One-way anova, *** p<0.001. N=6 images from 3 mice. (J) Record of body weight change during 0-42 dpi in PS5A1 GBM treatment. N=5 mice in each group.

Since most chemotherapy drugs are administered over multiple doses with intervals for tolerance and recovery, it is important to assess the feasibility of multiple BBTB openings for drug delivery. Taxol (generic name paclitaxel) is a microtubule-stabilizing drug approved by the FDA for the treatment of ovarian, breast, and lung cancer, as well as Kaposi’s sarcoma (*27*). Following the failure of Taxol to show efficacy in an early-phase clinical trial for GBM, further testing was abandoned. However, Taxol cannot pass through the BBB and this may in part account for the lack of clinical efficacy. To this end, we co-delivered AuNP-BV11 and fluorescent dye-labeled Taxol (Taxol Janelia Fluor 646) to PS5A1 GBM-bearing mice to investigate multiple BBTB openings during tumor treatment. **Fig. 2C** shows that the first cycle of optoBBTB at 14 dpi led to an increase in dye permeability in the tumor core and margin area, as did the second and third BBTB cycles (18 and 22 dpi). Each resulted in dye leakage primarily in the tumor core and margin regions. Notably, there was no evidence of fluorescent taxol leakage in the contralateral hemisphere which serves as an internal control and reconfirms the inability of this drug to pass through the normal BBB. Accordingly, for subsequent in vivo testing of Taxol in PS5A1 GBM model, we used a three-cycle treatment regimen.

### OptoBBTB enables delivering a non-BBB penetrant chemotherapy drug and improves therapeutic outcome for PS5A1 GEMM GBM model

To investigate the efficacy of optoBBTB for Taxol delivery in our GEMM, we first evaluated its efficacy and mechanism of action in vitro. Taxol is known to binding to the mitotic spindle apparatus, and disrupt chromosomal segregation which leads to mitotic catastrophe and cell death (*28, 29*). In vitro studies showed the internalization of Taxol Janelia Fluor 646 in PS5A1 glioma cells after 1-hour co-incubation. PS5A1glioma cells were sensitive to the Taxol with an IC50 value of 17 nM at 72 hours (**Fig. S4**). However, approximately 20% of tumor cells were resistant to Taxol, whether these cells represent an inherently resistant subpopulation or they acquire resistance following drug exposure, is not clear and is the subject of our ongoing work. Importantly, this observation underscores that our choice of PS5A1 glioma line mimics the clinical scenario of GBM heterogeneity which is thought to be a major mechanism of drug resistance. To further assess the treatment efficacy of optoBBTB in PS5A1 glioma-bearing mice, mice were randomly grouped at 14 dpi and treated with (1) vehicle, (2) free Taxol (i.v. administration, 12.5 mg/kg), (3) optoBBTB followed by vehicle delivery, and (4) optoBBTB followed by taxol delivery (i.v. administration, 12.5 mg/kg). The treatment regimen included 3 cycles delivered every 4 days from 14 to 22 dpi, and the treatment efficiency was evaluated at 42 dpi (**Fig. 3A**). The results show that optoBBTB greatly enhanced the delivery of fluorescent Taxol646 in the tumor core and margin compared to no optoBBTB treatment (**Fig. 3B-C**). These tumors showed no T1W contrast enhancement by MRI consistent with an intact BBB and minimal T2W hyperintensity. This observation is consistent with the clinical scenario where GBM cells are undetectable to convention MR sequences (*30*). To unequivocally verify presence of glioma cells, we collected all the tumor-containing brain slices and analyzed the tumor volume by GFP fluorescence. Remarkably, optoBBTB+taxol yielded the smallest tumor volume (68.09±20.77 mm^3^), a 5 to 7-fold reduction when compared to vehicle (473.88±37.56 mm^3^), Taxol (442.11±134.83 mm^3^), and optoBBTB+vehicle (334.51±123.83 mm^3^, **Fig. 3D**). Moreover, optoBBTB+taxol delivery significantly increased the survival by 50%, from 40 days to 60 days (**Fig. 3E**), as a result of a marked inhibition of tumor growth (**Fig. 3F, 3G top, Fig. S5**). Furthermore, Ki67 staining and cell apoptosis analysis showed that treatment with optoBBTB and Taxol decreased cell proliferation, increased cellular apoptosis in tumors compared to other groups, and prolonged animal survival by 50% from 40 to 60 days (**Fig. 3G-I**). There was no significant difference in body weight at 42 dpi (**Fig. 3J**), indicating that optoBBTB plus Taxol did not induce significant systematic toxicity. These results supported that optoBBTB slowed tumor progression and extended survival in infiltrative GBM-bearing mice.

### 73C glioma model shows heterogeneous BBTB integrity during disease progression

Next, we evaluated the growth and BBTB permeability using a genetically engineered mouse glioma model (Braf^V600E^, p53^-/-^, Pten^-/-^; see supplementary information for details), which is characterized by aggressive expanding mass with limited diffuse single-cell infiltration and a hypoxic core region reminiscent of clinical tumor core regions (*31*). These cells were derived from conditional multi-allele (LSL.Braf^V600E^, Pten^f/f^, p53^f/f^) primary astrocytes and infected ex vivo with an adeno-Cre virus to generate a primary transformed cell line (referred to as 73C). These cells were also engineered to express GFP. The BBTB integrity during 73C glioma progression was analyzed by i.v. injection of fluorescent dyes EZ-link biotin (660 Da) and Evans blue (66 kDa, albumin-bound) at 7, 14, and 21 days post 73C cell injection (dpi), respectively. **Fig. 4A and B** show that at 7 dpi, both low and high molecular weight fluorescent dyes were limited to the microvascular lumen in the tumor core region, at the margins of the expanding mass which interface with the normal brain parenchyma, and at the contralateral hemisphere. This result suggests an intact BBTB at 7 dpi. In contrast, at 14 and 21 dpi, following the rapid expansion of the tumor mass, there was clear evidence of dye extravasation in the tumor core regions but absent at the tumor margins. These observations suggest that the tumor core region is perfused by a microvasculature with compromised BBB integrity while the tumor margin regions which interface with the surrounding brain parenchyma are able to co-opt the normal brain capillary network.

**Fig. 4.**
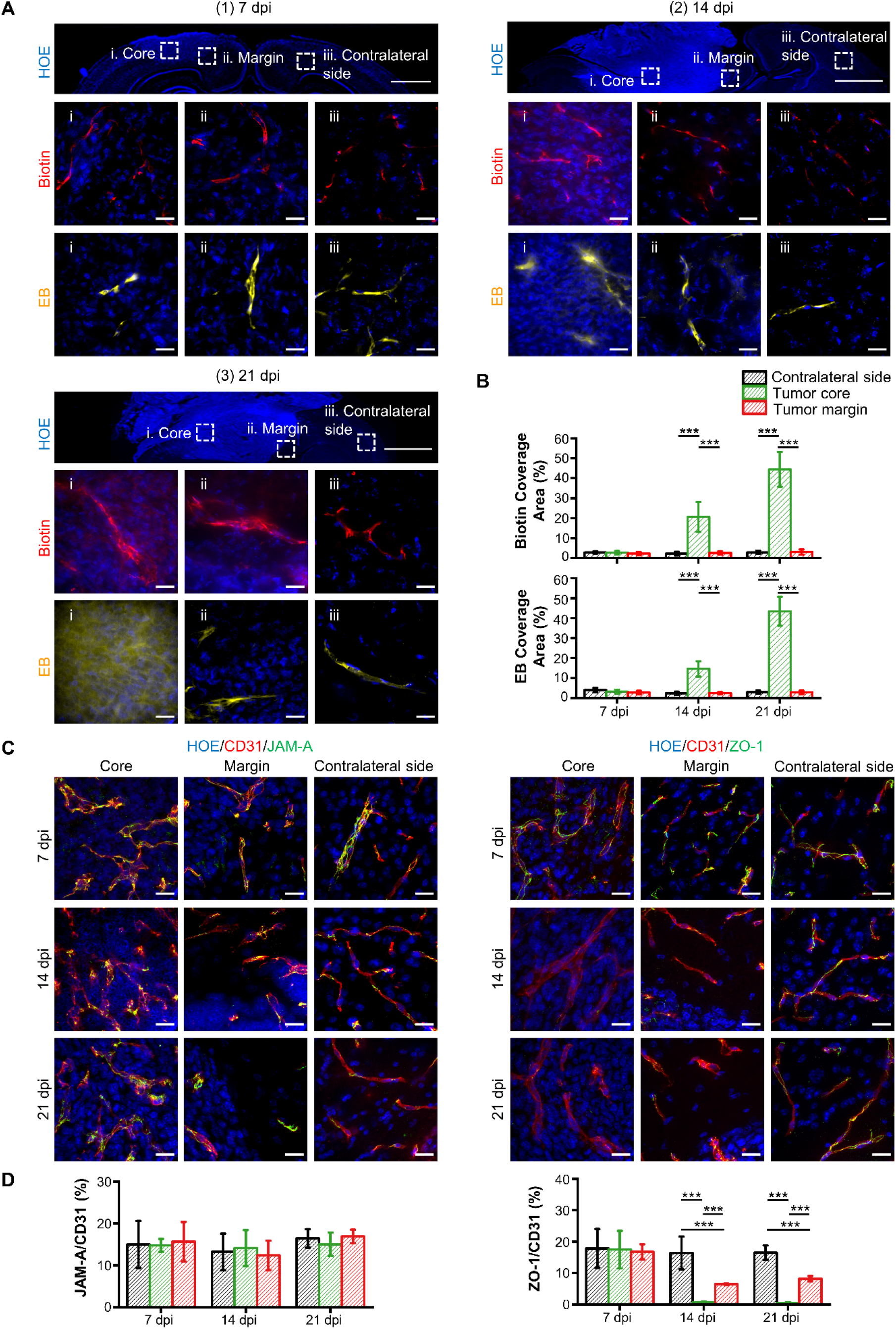
73C GBM model shows heterogeneous loss of BBTB integrity during disease progression. (A) BBTB permeability analysis by using EZ-link biotin (Biotin, red) and Evans blue (EB, yellow) at 7 dpi, 14 dpi, and 21 dpi (dpi: days post-injection). The cell nuclei were indicated by Hoechst staining (HOE, blue). The ROIs selected are (i) tumor core, (ii) tumor margin, and (iii) contralateral side with no tumor. The scale bar represents 1 mm in the upper panel and 20 µm in the middle and bottom panels. (B) The quantification of biotin and Evans blue coverage by area fraction. One-way Anova, ***p<0.001, N=15 images from 3 mice. (C) IHC staining of junctional protein JAM-A and ZO-1 at the selected time points. The blood vessels were stained with CD31 (red), and the cell nuclei were indicated by Hoechst staining (HOE, blue). The scale bar represents 20 µm. (D) Quantification of JAM-A and ZO-1 coverage on the blood vessel by area fraction. One-way Anova, ***p<0.001, N=15 images from 3 mice.

It is well established that tumor core regions under hypoxic stress activate proangiogenic signals leading to the formation of immature and dysfunctional network (*32*). To test this hypothesis, we further evaluate the immunostaining of TJ-associated proteins, Claudin-5, ZO-1, VE-Cadherin, and JAM-A with CD31-labeled endothelial cells. Notably, CD31-labeled microvascular density significantly increased in the tumor core compared to the tumor margin and contralateral brain region (indicated by CD31+, P<0.001, **Fig. S6A**). In contrast, i.v. injection of tomato lectin to label perfused vessels showed a marked increase in the ratio of cell nuclei labeled with Hoechst dye (HOE) to the blood vessels (HOE/lectin) in the tumor, compared to the healthy contralateral side (**Fig. S6B,** top). However, there was no significant difference in this ratio with IHC staining of blood vessels using CD31 (HOE/CD31, **Fig. S6B**, bottom). These results suggest that the tumor core region contains a significant fraction of either nascent vessels which have yet to form a functional conduit and/or have formed non-functional end-vessels which are not perfused. IHC staining of junctional proteins showed that the immunofluorescence of Claudin-5, VE-Cadherin, and JAM-A persisted during 7-21 dpi at both tumor core and margin (**Fig. S6C,** **Fig. 4C, left**). However, there was a significantly lower level of ZO-1expression at the tumor core at 14 and 21 dpi (**Fig. 4C, right**), consistent with the observation in human GBM (**Fig. S1**). Further quantification analysis of the area fraction ratio of protein over blood vessel (CD31) suggested that the relative protein coverage ratio for Claudin-5, VE-Cadherin, and JAM-A was comparable at the tumor core, margin, and contralateral side during 7-21 dpi, while their expression in the tumor area was elevated compared to contralateral side due to the increased blood vessel coverage (**Fig. S6C,** **Fig. 4D, left**). Although no apparent change in the ZO-1/CD31 was observed at 7 dpi, there is a significant decrease at the tumor core and margin at 14 and 21 dpi (**Fig. 4D, right**). We speculated that the BBTB disruption in the tumor core during disease progression was partially due to the loss of ZO-1 coverage on immature newly formed vessels. Taken together, our spatiotemporal analysis of tumor vasculature suggests that the 73C orthotopic GBM model features robust angiogenesis and immature dysfunctional tumor associate vessels and is reminiscent of clinical tumors. Therefore, it is a suitable surrogate for assessing drug delivery strategies.

### OptoBBTB improves drug delivery and therapeutic outcomes in 73C GEMM GBM model

We subsequently investigated the efficacy of optoBBTB in 73C GBM model. The overexpression of JAM-A in the tumor area makes the nanoparticles attractive for enhanced GBM accumulation. Inductively coupled plasma mass spectrometry (ICP-MS) analysis shows >80% increase of AuNP-BV11 accumulation in the tumor compared to normal brain tissue (0.102±0.016 versus 0.188± 0.025 %ID/g in normal brain and tumor, respectively, **Fig. 5A**). Biodistribution of the injected particles in other organs is consistent as previously reported (**Fig. S2E**) (*23*). We further characterized the optoBBTB in the 73C GBM model (7 dpi) to obtain the optimal opening efficiency after single-pulse ps laser stimulation by assessing a variety of nanoparticle injection doses and laser fluence (**Fig. S7A, Table S2**). The highest BBTB opening level was achieved by injecting 36 µg/g AuNP-BV11 followed by 40 mJ/cm^2^ laser excitation. Since the BBTB in the tumor core and the margin remained intact at 7 dpi, optoBBTB significantly improved the delivery of both small molecules (EZ-link biotin, 660 Da) and large molecules (Evans blue, 66 kDa) after i.v. injection (**Fig. 5B**). The BBTB opening was gradually recovered in 72 hours, and no dye leakage into the brain was observed (**Fig. S7B**). Similarly, a three-cycle treatment regimen was used for the 73C GBM model (**Fig. 5C**), while no optoBBTB modulation at 12 dpi showed no obvious dye leakage into the tumor (**Fig. S7C**).

**Fig. 5.**
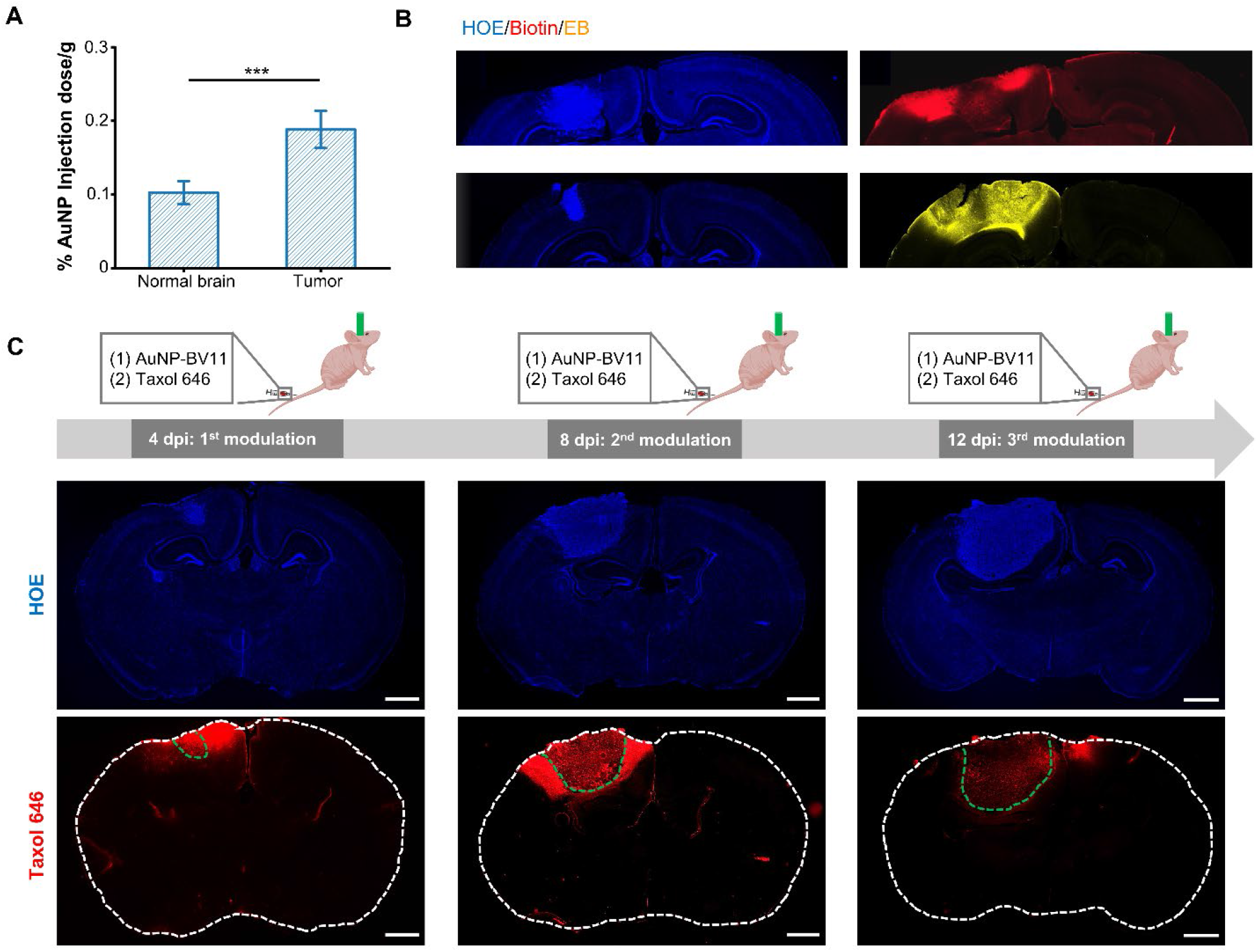
OptoBBTB improves drug penetration in 73C GBM model. (A) The analysis of AuNP-BV11 accumulation in 73C tumor-bearing mice. Student’s t-test, n=3 mice, ** p<0.01. (B) optoBBTB allowed the delivery of small molecule EZ-link biotin (660 Da) and large molecule Evans blue (66 kDa, albumin-bound) to the tumor. (C) Multiple BBTB modulations in 73C GBM model at 4, 8, and 12 dpi. AuNP-BV11 and taxol646 were injected intravenously. The cell nuclei were indicated by Hoechst staining (blue), and taxol646 was indicated by red. The scale bar represents 1 mm.

Then we analyzed the cellular uptake and the cellular toxicity of Taxol in 73C glioma cells. After co-incubating 73C glioma cells and Taxol Janelia Fluor 646 (3 µM) for 1 hour, Taxol was accumulated in the microtubules enriched cytoplasm (**Fig. S8A**), consistent with previous report (*33*). Next, 73C glioma cells were co-cultured with taxol over a 3-log fold concentration range (0-1000 nM for 72 hours and cell viability was determined using WST-1 assay. Our results showed that while Taxol was highly potent against 73C glioma cells, with an IC50 value of 37 nM (**Fig. S8B**) measured by conventional dose-response measures, however, its maximum effect seen at ∼70 nM, with approximately 35% of the remained viable despite increasing drug concentration to 1000 nM. This suggests that either a surviving fraction of 73C cells rapidly acquire resistance to Taxol and remain viable over the duration of the assay. Alternatively, since the 73C glioma cells represent a polyclonal population, it may include a subpopulation of inherently Taxol resistant cells. This information is critical for interpreting in vivo results with optoBBTB and closely mimics the increasingly recognized clinical scenario of tumor cell heterogeneity with subpopulations of drug-resistant tumor cells.

Next, we characterized the optoBBTB in tumor treatment using 73C glioma-bearing mice. The initial tumor volume was measured by magnetic resonance imaging (MRI) at 3 dpi, and the mice were randomly grouped and treated with (1) vehicle, (2) taxol administration (12.5 mg/kg), (3) optoBBTB followed by vehicle administration, and (4) optoBBTB followed by taxol administration (12.5mg/kg). We performed three treatments on a 4-day interval (covering dpi 4-12). At 15 dpi, the tumor volume was measured by MRI, and the brains were harvested for histological analysis (**Fig. 6A**). MRI T2-weighted scan was used to measure the tumor volume since T1-weighted gadolinium (Gd) enhancement showed low signal intensity, probably due to the intact BBTB at the early tumor stage (e.g., 3 dpi, **Fig. S9A**). **Fig. 6B-C** shows that a single dose of optoBBTB enhanced the delivery of Taxol to the tumor core and margin compared to no optoBBTB treatment. This enhanced Taxol delivery produced a statistically significant difference in slowing the tumor progression and increasing survival (**Fig. 6D-E** and **Fig. S9B**). Tumor volume analysis by MRI showed that optoBBTB+taxol delivery resulted in a 2.2 to 2.6-fold volume reduction in the tumor (35.68±7.32 mm^3^) compared to vehicle (92.75±19.1 mm^3^), Taxol (77.08±8.11 mm^3^), and optoBBTB+vehicle delivery (86.96±13.79 mm^3^) by MRI (**Fig. 6D, Fig. S9C**). The smallest tumor size in the group of optoBBTB+taxol was also confirmed by fluorescent imaging (**Fig. 6F**) and Hoechst staining of cell nuclei (**Fig. 6G**, top). (*23*). Consistent with the smaller tumor volume seen at 15dpi, overall median survival also increased from 18 days to 24 days by optoBBTB+taxol (**Fig. 6E**). Despite the modest 33% increase in survival duration, it is important to note that 73C glioma model shows no improvement in overall survival (OS) when treated with temozolomide (50 mg/kg/day), which is the FDA approved first-line therapy for gliomas (data not shown). Also, its efficacy is limited by the emergence of the taxol resistant subpopulation evident in our in vitro studies (see above and **Fig. S8B**). Immunohistological analysis of the optoBBTB+taxol tumors showed a marked decrease in proliferation (Ki67 positive cells) as well as an increase in cell death (Tunnel positive cells) relative to the other cohorts (**Fig. 6G-I**, middle, bottom). These data suggest that following BBTB disruption, Taxol can induce both cell cycle arrest and cell death. No significant difference in body weight was observed by the end of three treatments (**Fig. 6J**). Taken together, our data show that optoBBTB is able to significantly enhance the delivery of Taxol to a glioma model with a heterogeneous pattern of BBB integrity, and sufficient to induce tumor cell-cycle arrest and as well as improve overall survival.

**Fig. 6.**
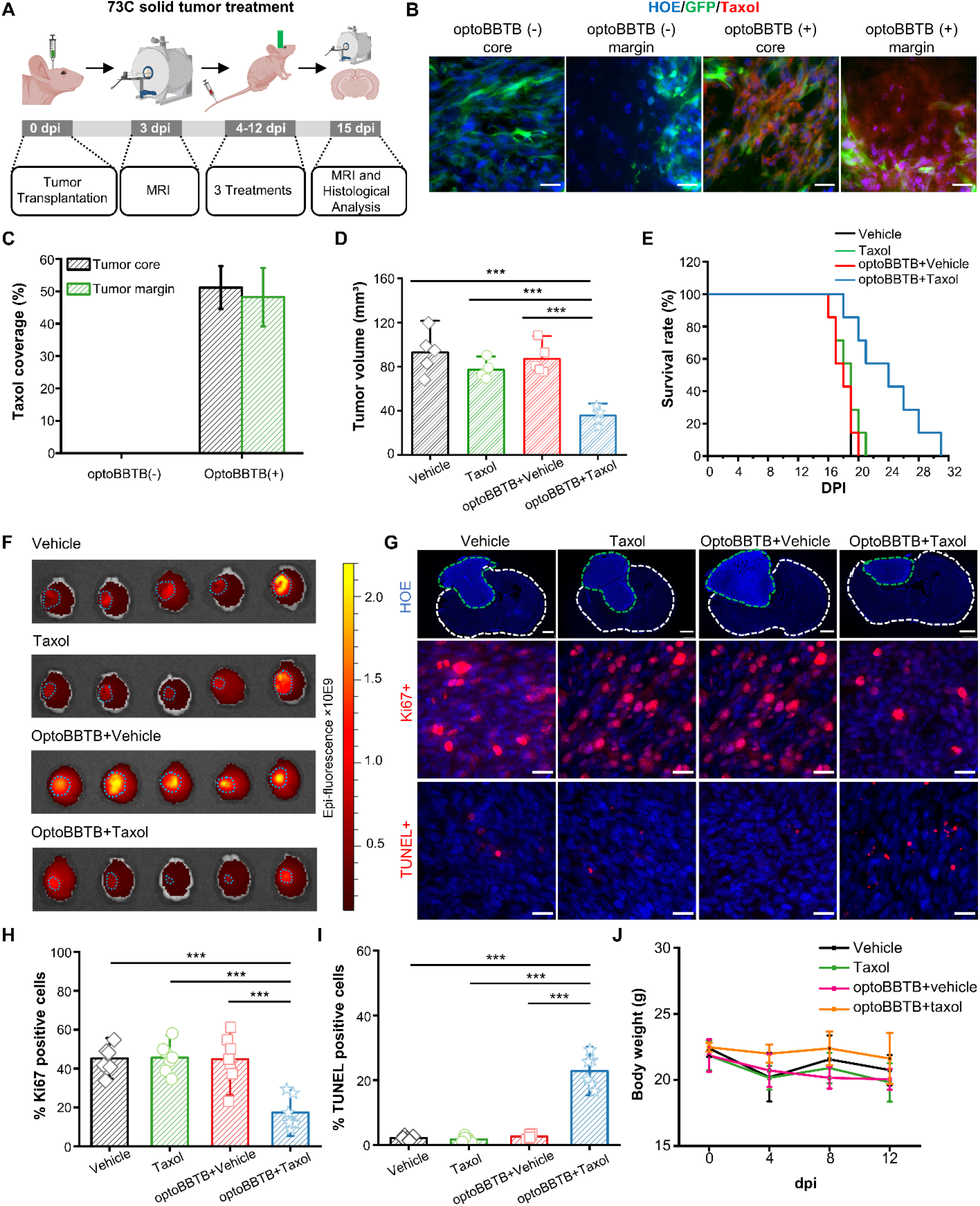
OptoBBTB improves therapeutic outcomes in the 73C GBM model. (A) schematic illustration of the treatment process. (B) OptoBBTB facilitated the delivery of fluorescent Taxol to tumors. The scale bar represents 20 µm. (C) Quantification of taxol delivery by fluorescent area fraction. (D) The analysis of tumor volume was measured by MRI at 15 dpi. One-way Anova, ***p<0.001, n=5 mice in each group. (E) Kaplan-Meier survival analysis, N=7 mice in each group. (F) Fluorescent imaging of tumor size at 15 dpi. (G) Top: Tumor size indicated by fluorescent imaging at 15 dpi using Hoechst staining (HOE). The scale bar represents 1mm. Middle: Ki67 staining showing cell proliferation, scale bar represents 20 µm. Bottom: TUNEL staining indicates cell apoptosis. Scale bar=20 µm. (H-I) Quantification of Ki67 staining and TUNEL staining after 73C GBM treatment. One-way anova, *** p<0.001. N=6 images from 3 mice. (J) Record of body weight change during 0-12 dpi in PS5A1 GBM treatment. N=5 mice in each group.

## Discussion

This devastating prognosis for GBM patients underlines the critical need to develop new therapies. Since one of the obstacles in GBM treatment is the existence of BBTB, inducing a temporary window to elevate BBTB permeability will enhance drug delivery to the tumor site. In this study, we showed that the optoBBTB technique reversibly increases BBTB permeability in both infiltrative and solid tumors. Importantly, these nanoparticles were selectively accumulated in the 73C glioma model due to the JAM-A overexpression compared to the healthy tissue. We showed that the combination of optoBBTB and Taxol delivery significantly suppressed tumor growth and prolonged the survival of tumor-bearing mice without causing adverse effects. These results demonstrated that optoBBTB is a selective, safe and reversible technique to overcome the BBTB.

Intercellular TJs represent a formidable barrier against paracellular drug delivery at the BBB (*14, 34*). Several approaches have been developed to modulate the TJ to enhance the delivery across the BBB, including co-administration of siRNA against claudin-5 and occludin, as well as exploiting claudins or cadherin inhibitory peptides (*35–38*). However, impediments to successful translation of these approaches were reported, such as a lack of a robust delivery system in human, poor targeting efficacy, or a lack of site-specificity (*39, 40*). Efforts have also been devoted to modulating tumor TJs, while the efficacy and safety concerns required further evaluation (*14*). We demonstrated that JAM-A could be targeted and modulated to increase the BBTB permeability, owning advantages such as non-invasive and high spatiotemporal resolution. According to our previous study, the increase in permeability is partially due to the increased paracellular diffusion and the widened TJ cleft after picosecond laser excitation of AuNP-BV11 (*23*). Moreover, we also observed overexpression of Claudin-5 and VE-Cadherin in 73C glioma in addition to JAM-A, which may serve as therapeutic targets in certain types of GBM.

One crucial contributing factor to the poor prognosis of GBM is the use of preclinical models that fail to fully recapitulate GBM pathophysiology, preventing efficient translation from the lab into successful therapies in the clinic (*20*). Xenograft mouse tumor models have been widely exploited in GBM treatment research in the past decades. However, when translating the results from in vitro to in vivo, one should consider the complex nature of the disease and the intrinsic limitation of these cell lines. For example, U87 and GL261 glioma cells have been used extensively (*41–43*). However, concerns have been rising recently, including the alteration of the genetic profile, interlaboratory variability, irrelevant gene mutation, and failure in recapitulating the immunogenicity of human GBM (*44–46*). Altogether, selecting appropriate in vitro and in vivo preclinical glioma models is critical to improving the efficacy of preclinical testing and ultimately making the discovery and translation of GBM therapies a more efficient process. To this end, we utilized two glioma cell lines (PS5A1 and 73C) to establish glioma models (*31, 47*). PS5A1 glioma showed an infiltrative growth pattern with vessel co-option development, and 73C glioma recapitulated GBM properties such as angiogenetic tumor core and intratumorally heterogeneity in terms of BBTB function and TJ composition. Moreover, they carried several critical mutations such as Pten, p53, LSLBraf^V600E^ and Ink4ab/Arf. The choice of these mutations was based on the high prevalence of p53 pathway inactivation and loss of Pten, while BRAF^V600E^, a rare mutation in adult GBM and common in the pediatric tumors, represents powerful co-activation of the phosphatidylinositol 3-kinase (PI3K)/AKT and mitogen-activated protein kinase (MAPK)/ERK signal transduction pathways. Both of these pathways are ubiquitously activated in GBM (*47*). Moreover, about 60% of GBMs show Ink4ab/Arf deletion or inactivation. These tumor cell lines allow us to assess the GBM therapy and interpret preclinical outcomes under the clinically relevant circumstance, which is fundamental for the transfer of novel technologies into clinical practice.

We noticed that the optoBBTB displayed a higher BBTB opening efficiency in the PS5A1 glioma model than in the 73C glioma model. Although we observed taxol delivery into both tumor core and margin after optoBBTB, suppressed tumor progression, and prolonged survival rate in the 73C glioma model, these effects were more predominant in PS5A1 glioma. We attempted to functionalize AuNP with other vasculature targets such as vascular endothelial growth factor 2 (VEGFR2) and transferrin receptor (TfR) that were overexpressed in 73C glioma. However, these nanoparticles didn’t improve BBTB opening efficiency compared to AuNP-BV11 (**Fig. S10**). Indeed, we observed poor vascular perfusion in the 73C glioma model (**Fig. S6)**. The lack of functional blood vessels may compromise the drug delivery efficacy in the tumor region (*48*). Moreover, blood vessels from angiogenesis may respond differently to the laser excitation than normal blood vessels, leading to lower BBTB disruption in the tumor core. Further investigation is needed to study how the normal and angiogenesis blood vessels respond to laser stimulation in terms of calcium influxes and cytoskeleton protein organization, which are key parameters in regulating BBB permeability (*49*). Another reason is that the depth of light penetration can be influenced by the cellular density of the tumor tissue (*50*). Different from the infiltrative growth of the PS5A1 glioma, the tumor cells are densely packed in the tumor core in 73C glioma, probably resulting in insufficient light penetration in this area. In this context, we speculate that optoBBTB can be used in different stages in GBM therapy, such as drug delivery to the angiogenesis GBM model, or the infiltrative margin after surgical removal of the tumor.

Although optoBBTB followed by Taxol delivery showed exciting outcomes in delaying tumor proliferation and prolonging survival by 33.3% and 50% in 73C and PS5A1 mouse tumor models, it didn’t result in a complete cure. We observed that in vitro treatment of PS5A1 and 73C glioma cells with Taxol for 72 hours resulted in 20-30% live cells (**Fig. S4 and S8**). The development of taxol tolerance could be attributed to several aspects. First, the overexpression of P-glycoprotein (P-gp), an ATP-binding cassette (ABC) drug efflux pump, on blood microvasculature and tumor cell membrane lowers intracellular drug concentration by accelerating drug efflux, making it a common mechanism of drug resistance (*51, 52*). P-gp can transport a wide range of molecules including Taxol, resulting in an insufficient drug amount to completely regress the tumor (*53*). Second, chemotherapy is most effective at killing cells that are rapidly dividing. Dormant tumor cells after optoBBTB and taxol delivery may have active drug resistance that prevents them from complete removal (*54*).

There are several considerations for further validation and translation of optoBBTB. First, improvement of the optoBBTB approach includes exploring near-infrared laser NIR-absorbing nanoparticles since the light in this region exhibits deeper tissue penetration to cover the needed tumor margin. Second, while light can be delivered transcranially in the mice brain, fiber delivery to the human brain is envisioned, especially after surgical removal of the primary tumor. Placing an optical fiber in the tumor surgical cavity would allow light delivery to the tumor margin.

In summary, this work provides a new strategy to modulate the BBTB permeability by manipulating its TJ component. OptoBBTB improved the delivery of Taxol to both clinically relevant solid and infiltrative tumors, suppressed tumor growth, and effectively extended survival. OptoBBTB has the potential for clinical translation to improve GBM therapy.

## Materials and methods

### Materials

Gold (III) chloride trihydrate, hydroquinone, sodium citrate tribasic dihydrate, endotoxin-free ultrapure water, Evans blue, EZ-link biotin, Dimethyl sulfoxide (DMSO), Tween 20, Triton X-100, Paclitaxel (Taxol), Cremophor® EL, sucrose, Epidermal growth factor (EGF) and Recombinant Human Fibroblast Growth Factor Basic (FGF2) Progesterone were purchased from Sigma-Aldrich. OPSS-PEG-SVA (3400 Da), and mPEG-thiol (1000 Da) were purchased from Laysan Bio, Inc. Endotoxin-free ultrapure water, Donkey serum, phosphate-buffered saline (sterile), saline (sterile), Hoechst stain (33342), borate buffer, paraformaldehyde (4% in PBS), gold reference standard solution, lectin Dylight 488/594/647, 20 kDa molecular weight cut off (MWCO) dialysis membrane, Gibco^™^ DMEM (high glucose, GlutaMAX^™^, DMEM), Dulbecco’s Modified Eagle’s Medium)/F12 50:50 Mix (DMEM/F12), fetal bovine serum (FBS), StemPro™ Accutase™ Cell Dissociation Reagent, trypsin-EDTA solution (0.25%), penicillin-streptomycin, Tissue-Plus^™^ O.C.T. compound, Invitrogen Fluoromount-G^™^ mounting medium, B-27^TM^ supplement, Insulin Transferrin solution and Cayman Chemical WST-1 Cell Proliferation Assay Kit were purchased from Thermo Fisher Scientific. All chemicals were analytical grade unless specified. Taxol Janelia Fluor^®^ 646 was purchased from Tocris Bioscience. ApopTag^®^ Red In Situ Apoptosis Detection Kit (S7165) was purchased from Millipore Sigma. Buprenorphine SR-LAB 5 mL (1 mg/mL) was purchased from ZooPharm.

Anti-JAM-A antibodies BV11 and BV12 (for IHC staining) were provided by Drs Elisabeth Dejana and Monica Giannotta at FIRC Institute of Molecular Oncology Foundation. Rat anti-CD31, goat anti-CD31, rat anti-transferrin (8D3), rat anti-VEGFR2 (BE0060), rabbit anti-Claudin-5 (341600), rabbit anti-ZO-1(402200), rabbit anti-VE-cadherin (361900), rabbit anti-ki67, donkey anti-goat IgG Alexa 488, donkey anti-rat IgG Alexa 594, donkey anti-rabbit IgG Alexa 594/647 were purchased from Fisher Scientific unless otherwise noted.

### Animals

The immunodeficient nude mice Foxn1nu (Nu/J, stock number 002019, 7 weeks old, female, 20-25 g) were ordered from Jackson Laboratories. Animal protocols were approved by Institutional Animal Care Use Committee (IACUC) of University of Texas at Dallas.

### Human GBM sample

Under an Institutional Review Board (IRB)-approved protocol, patients with brain tumors are approached by an attending neurosurgeon, or a clinical research coordinator at their initial visit to the University of Texas Southwestern Medical Center and asked to give consent to the collection of blood and residual tissue samples in the case of surgical resection. Patient recruitment and consent, as well as blood and tumor sampling, have been previously described (*55, 56*)

Fresh brain tumor samples were obtained from adult patients during their operative procedure after informed consent was obtained. Brain tumors were graded and tumor core and tumor margin were also defined by T2-weighted fluid-attenuated inversion recovery (FLAIR) Imaging of patient brain, at the Southwestern Medical Centre, University of Texas, by a neuropathologist according to World Health Organization guidelines.

Fresh tumor samples were fixed in formalin and 4% PFA, for paraffin embedding and Cryogel embedding, respectively. Five-micrometer thick formalin-fixed paraffin-embedded GBM samples were processed for H&E staining, Ten-micrometer thick PFA-fixed Cryogel-embedded samples were processed for immuno-fluorescence staining. During staining the slides were de-paraffinized and rehydrated by serial incubations in xylene (2X for 3 min), 100% ethanol (2X for 3 min), 95% ethanol (2X for 3 min), 70% ethanol (2X for 3 min) and PBS (4X for 3 min). To prevent sections from folding or falling, they were incubated in 10% NBF for 30 min. Following this, antigen retrieval was carried out by placing slides in 700 mL of Antigen retrieval buffer (10 mM Tris HCl, 1 mM EDTA, 10% glycerol, pH 9.0) at 95℃ for 25 min. The slides were then cooled to room temperature for 30 min and rinsed with PBS (2X for 2 min) before being blocked in 10% horse, donkey, or goat serum (depending on the species of the secondary antibodies) in 0.1% PBS-T for 1 h at room temperature. The slides were incubated with primary antibodies in 2% serum PBST overnight at 4 ℃. The next day, the slides were washed in PBS (3X for 10 min) and incubated with secondary antibodies for 1hr at room temperature. For fluorescence detection, fluorophore conjugated antibodies were used for secondary staining antibody of BBB marker protein, Rabbit anti ZOL1 (#617300, Invitrogen) and Rabbit anti-Occludin (#711500, Invitrogen) as primary antibody was used for immunofluorescent staining, as well as endothelial cell marker CD31 antibody (Rat anti CD31, #553370, BD Pharmingen), with general dilution factor 1:400 in PBS. Fluorescence conjugated secondary antibody was used accordingly. H&E staining was scanned on Nanozoomer 2.0HT (NDP), Hamamatsu Photonics K.K. (Japan) at 2X magnification. Immunofluorescence staining was scanned on Zeiss LSM 780 Confocal Microscope at 40x lens

### Glioma cell line and cell culture

73C glioma cells were obtained from Dr. Robert Bachoo’s laboratory, which were generated in primary astrocyte cultures from neonatal mice that carried conditional mutations for Pten^f/f^, p53^f/f^, and LSLBraf^V600E^ (*31*). 73C glioma cells were cultured in DMEM containing 10% fetal bovine serum (FBS) and 1% penicillin-Streptomycin.

PS5A1 glioma cells were obtained from Dr. Robert Bachoo’s laboratory. It is a highly invasive mouse glioma cell line which was derived from de novo glioma in the adult BL6 background conditional mouse (*Braf^V600Ef/+;^ Ink4ab/Arf ^f/f^; Pten^f/f^*) that was induced by intracranial injection of AAV5-GFAP-Cre-GFP. These cells were then infected with a Lenti-GFP and selected by puromycin with stable green fluorescent protein expression. PS5A1 cells were cultured as free-floating neurospheres in DMEM/F12 medium, 2% of B-27, 20 ng/mL of EGF, and 20 ng/mL of FGF2, 20 ng/mL progesterone and 1% insulin transferrin solution.

#### In vitro dose responses

To test the in vitro dose-response of glioma cells’ interaction with Taxol, 3000 cells in 100 µL of cell culture medium were seeded in each well of the 96 well plate. A day later, 100 µL of taxol solution (0-1000 nM) was added to each well, and the solution was gently mixed and incubated in a CO_2_ incubator (37 ℃) for 72 hrs. Then the cell viability was measured by using WST-1 assay and reading the absorbance at 450 nm. The effect of the vehicle (Cremophor® EL and ethanol) was excluded by incubating the cells with the vehicle and the absorbance at 450 nm was substituted from the reading.

#### In vitro cellular uptake of fluorescent Taxol

Glioma cells were seeded in a glass-bottom culture dish (35 mm petri dish with 10 mm microwell) with a density of 70000 cells/cm^2^. One day later, Taxol Janelia Fluor^®^ 646 was added to the dish to achieve a final concentration of 3 µM, and the cells were incubated for 1 hour. Then cells were washed and fixed with 4% PFA for 10 min. The cell nuclei were stained with Hoechst staining for 10 min. Finally, the cells were washed 3 times with PBS before imaging.

### Glioma cell transplantation

All glioma cells used for transplantation were passaged two days before transplantation to reach 70–80% confluence. The glioma cells were dissociated and resuspended in Hank’s Balanced Salt Solution (without Ca^2+^, Mg^2+^, HBSS). Cells were counted and a suspension was prepared with a density of 2×10^5^/µL.

Buprenorphine (1 mg/kg) was given to the mice subcutaneously before surgery. 100 nL of 73C glioma cell suspension or 360 nL of PS5A1 glioma cell suspension in HBSS was constantly injected into the mouse cortex (-1 mm, -1 mm, 0.5 mm) using a nanoinjector (World Precision Instrument) equipped with a glass micropipette. The glass micropipette has a 50 µm tip and was generated by a Micropipette Puller (P-97, Sutter Instrument Co.). After injection, a thin layer of glue was applied over the exposed skull and the surrounding skin, and a layer of body double (**Body Double™ Standard Set, Smooth-On, Inc.)** was applied to the skull for protection. The mice were housed for 3 days (73C) to 2 weeks (PS5A1) to allow glioma growth.

### Dye permeability experiment

To test the dye permeability in glioma-bearing mice, EZ-link biotin (660 Da, 2 mg/mL) and Evans Blue (66 KDa/albumin-bound, 2% w/v) were intravenously injected into two mice at 7-, 14- and 21-days post injection (dpi) after glioma cell transplantation for 73C glioma bearing mice, and (2) 2, 4 and 6 weeks after glioma cell transplantation for PS5A1 glioma bearing mice. 30 min after the dye injection, the mice were sacrificed by cervical dislocation, and the brains were extracted and post-fixed in 4% (w/v) paraformaldehyde (PFA) at 4 °C for 24 hrs. The brains were then snap-frozen on dry ice and cut to 20 µm thick coronal slices using a cryostat, and stained with cy3-streptavidin to detect biotin and Hoechst solution to label nuclei before imaging.

### Immunohistochemical (IHC) staining

To immunostain vascular biomarker (CD31) and junctional proteins (i.e. Claudin-5, ZO-1, VE-cadherin, and JAM-A), the mice brains were snap-frozen on dry ice once quickly removed from the skull and cut to 20 μm thick coronal slices on a cryostat. The brain slices were fixed for 10 min using ice-cold methanol at -20 °C. Blocking solution (5% normal donkey serum, 0.1% Triton X-100 in PBS) was applied to the tissue for 1 hour at room temperature. After washing, the sections were first incubated overnight at 4 °C with the following primary antibodies (dilution ratio was 1:500): (1) rat anti-CD31, rabbit anti-Claudin-5, rabbit anti-ZO-1, rabbit anti-VE-cadherin, or goat anti-CD31 and rat anti-JAM-A (BV12).

To immune stain transferrin receptor and vascular endothelial growth factor receptor 2 (VEGFR2), or to detect the cell proliferation after treatment, the mice were perfused with PBS and 4% PFA. The brains were extracted and post-fixed in 4% PFA overnight, then cut into 20 μm thick coronal slices on a cryostat. The slices were incubated in applied to the tissue for 1 hour at room temperature and then incubated with primary antibodies (1) goat anti-CD31, rat anti-Transferrin (8D3), rat anti-VEGFR2, or (2) rabbit anti-ki67 with a dilution of 1:500.

In all cases, the sections were then incubated with secondary antibodies (dilution ratio was 1:500) for 2 hrs at room temperature, followed by incubation with Hoechst solution (dilution ratio was 1:2000). The slides were washed with PBS and mounted with Fluoromount-G mounting medium for imaging.

#### The preparation of brain vascular-targeting gold nanoparticles

The method to prepare brain vascular-targeting gold nanoparticles (AuNPs-BV11) was adapted from our previously reported method (*23*). AuNP-TfR and AuNP-VEGFR2 were prepared using a similar approach. The concentration, size distribution, and morphology of the nanoparticles were analyzed using Ultraviolet-Visible Spectroscopy (UV-Vis), Dynamic Light Scattering (DLS), and Transmission Electron Microscopy (TEM).

#### Inductively coupled plasma mass spectrometry (ICP-MS)

ICP-MS was used to determine the biodistribution of AuNP-BV11 (37 µg/g) after intravenous (i.v.) injection to the 73C glioma-bearing mice at 7 dpi. 1 hour after nanoparticle injection, the mice were perfused with ice-cold PBS and the main organs were collected. The tissue was then digested in fresh aqua regia until the tissue was fully dissolved. Then the solution was centrifuged at 5000 g for 10 min, and the supernatant was collected and diluted with ultrapure water for ICP-MS analysis (Agilent 7900).

#### OptoBBTB optimization in glioma models

To optimize the BBTB modulation to achieve the highest opening efficiency, different nanoparticle doses and laser fluence conditions were tested with 73C and PS5A1 glioma-bearing mice as shown in Table S1 and S2. Evans blue dye (2% in PBS, 100 µL) and EZ-link biotin (2 mg/mL, 100 µL) was injected to visualize the BBTB modulation. 30 min after the laser excitation, the mice were perfused with PBS and 4% PFA and the brains were extracted and post-fixed with PFA overnight. The brains were then frozen on dry ice and cut into 20 µm thick slices using a cryostat.

#### Analysis of the BBTB opening window after laser excitation of AuNP-BV11

AuNP-BV11 (37 µg/g or 18.5 µg/g) was i.v. injected to 73C and PS5A1 tumor-bearing mice at 7 and 14 dpi, respectively (n=3 mice for each group). The mice received a ps laser pulse (40 mJ/cm^2^) after 1 hour. Then EZ-link biotin was injected at 2 min, 1 day, and 3 days after laser excitation. 30 min later after the injection, the brains were extracted and frozen on dry ice and cut into 20 µm thick slices using a cryostat. The brain slices were incubated with Cy3-streptavidin to detect biotin and Hoechst solution to stain cell nuclei.

#### Taxol delivery to the tumor after optoBBTB

AuNP-BV11 (37 µg/g and 18.5 µg/g for 73C and PS5A1 glioma bearing mice, respectively) was i.v. injected to a tumor-bearing mouse (7 and 14 dpi, respectively). 1 hour later, Taxol Janelia Fluor^®^ 646 (12.5 mg/kg) was i.v. injected, followed by applying a single pulse of ps laser (40 mJ/cm^2^). 6 hours after laser excitation, the mouse was perfused with PBS and 4% PFA, and the brain was extracted and then post-fixed in 4% PFA overnight. The brain was frozen on dry ice and cut into 20 µm thick slices using a crystat. The slices were stained with Hoechst to label the nuclei.

### optoBBTB for glioma treatment

For 73C glioma-bearing mice, the treatment was started on 4 dpi. The mice were randomly divided into four groups: (1) vehicle control; (2) free taxol control (12.5 mg/kg); (3) optoBBTB+vehicle; (4) optoBBTB+taxol (12.5 mg/kg) with 5 animals in each group. Specifically, Taxol was dissolved in a mixer of Cremophor EL: absolute ethanol (1:1 v/v, as the vehicle) to 6 mg/mL, and then diluted to 2 mg/mL with saline. To modulate the BBTB, 37 µg/g of the AuNP-BV11 was administrated by intravenous (i.v.) injection into the tumor-bearing mouse. 1 hour later, the mice received either vehicle or Taxol via i.v. injection. Then the mice received picosecond-laser excitation (40 mJ/cm^2^, 1 pulse, pulse duration was 28 ps, and beam size was 6 mm) in the tumor area. The treatment was repeated every four days, three treatments were conducted between 4-12 dpi. On 3 and 15 dpi, the tumor size at the start point and endpoint was measured by MRI (T2-Weighted scan).

The treatment for PS5A1 glioma-bearing mice was similar but started at 14 dpi. To modulate the BBTB, 18.5 µg/g of the AuNP-BV11 was i.v. injected and the laser fluence was 40 mJ/cm^2^. The treatment was repeated every four days and repeated 3 times. The mice were sacrificed at 42 dpi to analyze the tumor size.

To obtain the survival rate, similar treatment groups were used for 73C and PS5A1 glioma-bearing mice, with 7 mice in each group.

#### Magnetic Resonance Imaging (MRI)

73C glioma was visualized using a preclinical BioSpec 3T MRI (Bruker, Billerica, MA, USA) with a mouse head coil (Item RF Res 128 1H 064/023 QSN TF, Model 1P T167055, Serial S0013). T2-weighted scans were performed using a T2 RARE sequence with an echo time of 60 ms, repetition time of 2725.738 ms, an echo spacing of 15 ms, a rare factor of 10, 8 averages, 1 repetition, a slice thickness of 0.5 mm, a field of view of 20 by 20 mm, and a matrix size of 192 by 192. The number of slices ranged from 12 to 20 to cover the entire brain of each mouse. Images were exported to Digital Imaging and Communications in Medicine (DICOMs) and Fiji/Image-J for further analysis.

#### Fluorescent Microscopy, Electron Microscopy, and imaging analysis

All the fluorescent images were taken with Olympus FV3000RS confocal laser scanning microscope, Olympus SD-OSR spinning disk super resolution microscope, and Olympus VS120 virtual slide microscope. The transmission electron microscopy (TEM) images were taken using a JEOL JEM-2010 microscope.

For the IHC staining, the images were analyzed by Fiji/ImageJ. To study the changes in junctional proteins, the area fraction of Claudin-5, ZO-1, VE-cadherin, and JAMA-A was obtained and normalized by CD31 indicating cerebral vessel. Vasculature density was analyzed by area fraction of CD31.

To obtain the tumor volume after treatment in PS5A1 glioma-bearing mice, brain slices were imaged with Olympus VS120 virtual slide microscope. The total area with tumor GFP fluorescent was determined by selecting the optimal threshold and analyzed with Fiji/Image-J. The tumor volume was calculated by area×thickness.

#### In situ cell apoptosis detection

The cell apoptosis analysis was performed at 15 dpi (3 days after the 3^rd^ treatment) or 42 dpi (20 days after the 3^rd^ treatment) for 73C and PS5A1 glioma-bearing mice, respectively. The mice were perfused with PBS and PFA, and the mice’s brains were harvested and post fixed in PFA overnight. Then the brains were snap-freeze on dry ice. 30 µm thick brain slices were cut on a cryostat and the cell apoptosis detection was performed with ApopTag^®^ Red In Situ Apoptosis Detection Kit following the manufacturer’s protocol. Briefly, the slides were rinsed in two changes of PBS (5 min each), and post-fixed in pre-cooled ethanol: acetic acid 2:1 for 5 min at -20 ℃, followed by rinsing twice in PBS. Then equilibration buffer (75 µL/5 cm^2^) was applied to the specimen and incubated for 1 min at room temperature. Then the excess liquid was removed, and the TdT enzyme was applied to the samples (55 µL/5 cm^2^), followed by incubation at 37 ℃ for 1 hour. Subsequently, stop/wash buffer was added to the samples and incubated for 10 min at room temperature. The samples were washed three times in PBS, and anti-digoxigenin conjugate rhodamine was applied to the specimen (65 µL/5 cm^2^) and incubated for 30 min at room temperature. The samples were then washed four times in PBS, stain the cell nuclei with Hoechst staining (1:2000), and mount with Fluoromount-G^™^ for imaging.

## Statistical analysis

All the data were plotted and analyzed with Origin 2020. The indication of each data dot, n values per group, and details of statistical testing are provided in the figure caption.

## Author contributions

Q.C. and Z.Q. conceived the original idea. Q.C. performed the experiment and analyzed the data. X.G. and V.V. performed the IHC staining and imaging on human GBM samples. X.L., H.X., H.F. and X.G. assisted in experiments and analyzed the data. R.M. and J.L. performed magnetic resonance imaging. M.G. and E.D. prepared the BV11 and BV12 antibodies. R.B. and Z.Q. supervised the project, analyzed the data, and discussed the results. Q.C. and Z. Q. wrote the paper. All authors revised the manuscript and have approved the final version.

## Acknowledgment

We thank Dr. Jonghae Youn for assistance with TEM imaging, Mr. Chen Xie for assistance with photographs and Dr. Haihang Ye for reviewing and revising the manuscript.

## Data availability

All data associated with this study are present in the paper or the Supplementary Materials.

