## Supplemental File for "Optical Modulation of Blood-Brain-Tumor Barrier Permeability Enhances Drug Delivery in Diverse Preclinical Glioblastoma Models"

**This PDF file includes:**

Fig. S1-S10

Table S1-S2

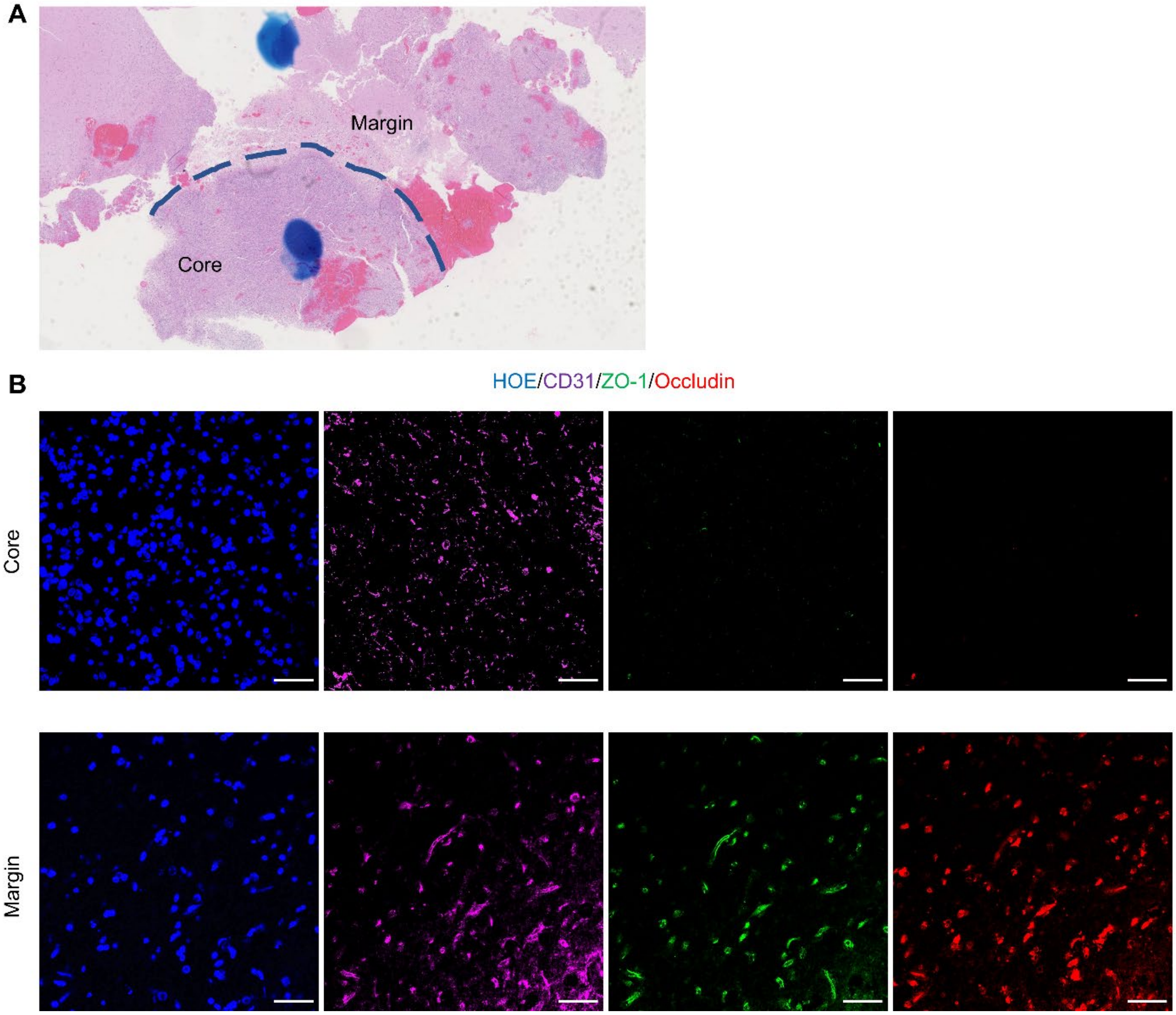


**Fig. S1. Human GBM shows core and margin regions with distinct tight junction protein expression.** (A) H&E staining showing the histopathological characteristics of human GBM. (B) IHC staining of key junctional proteins in tumor core and margin shows loss of ZO-1 and occludin expression in the core but not in the margin. Scale bar=50 μm.

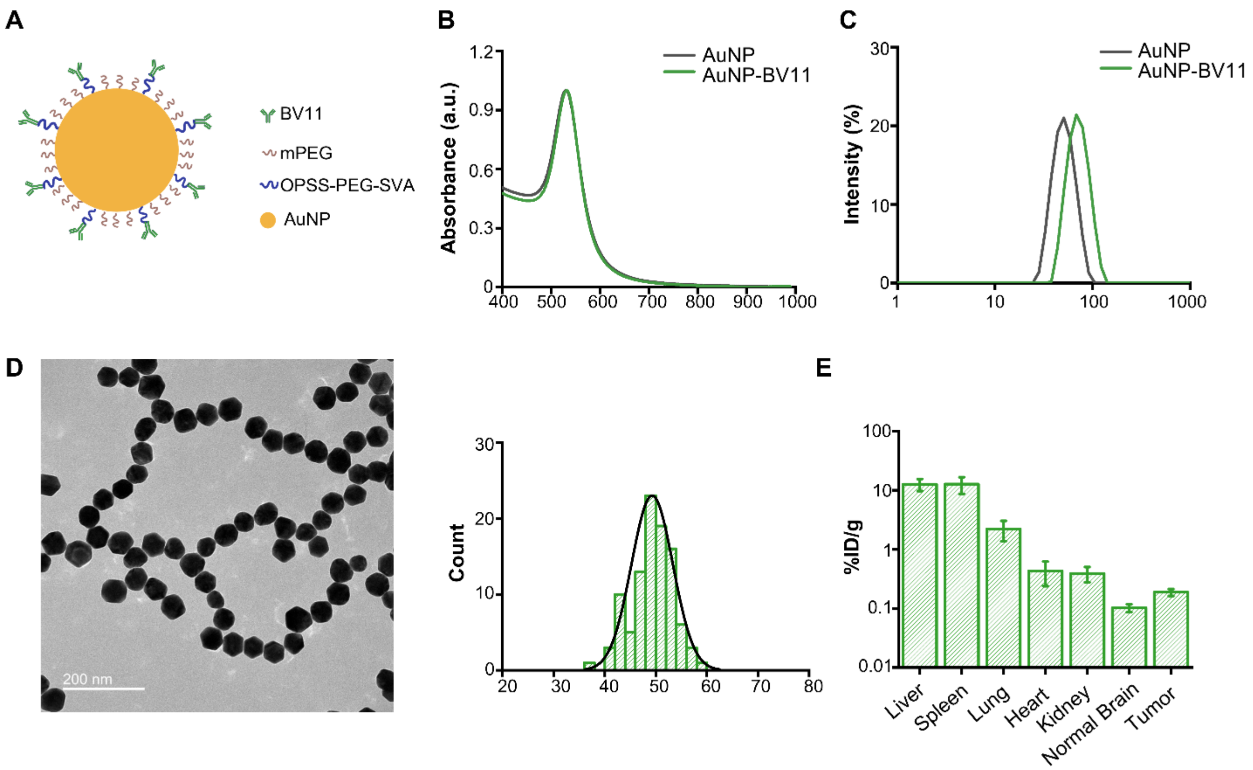


**Fig. S2. Characterization of AuNP-BV11 and its biodistribution in tumor-bearing mice.** (A) The surface functionalization of AuNP-BV11. (B) The LSPR peak is characterized by UV-Vis-NIR spectroscopy. (C) The size distribution is characterized by Dynamic Light Scattering (DLS). (D) The morphology and size distribution of the nanoparticles are characterized by TEM. (E) The biodistribution of AuNP-BV11 in 73C tumor-bearing mice. N=3 mice, data were expressed as mean±SD.

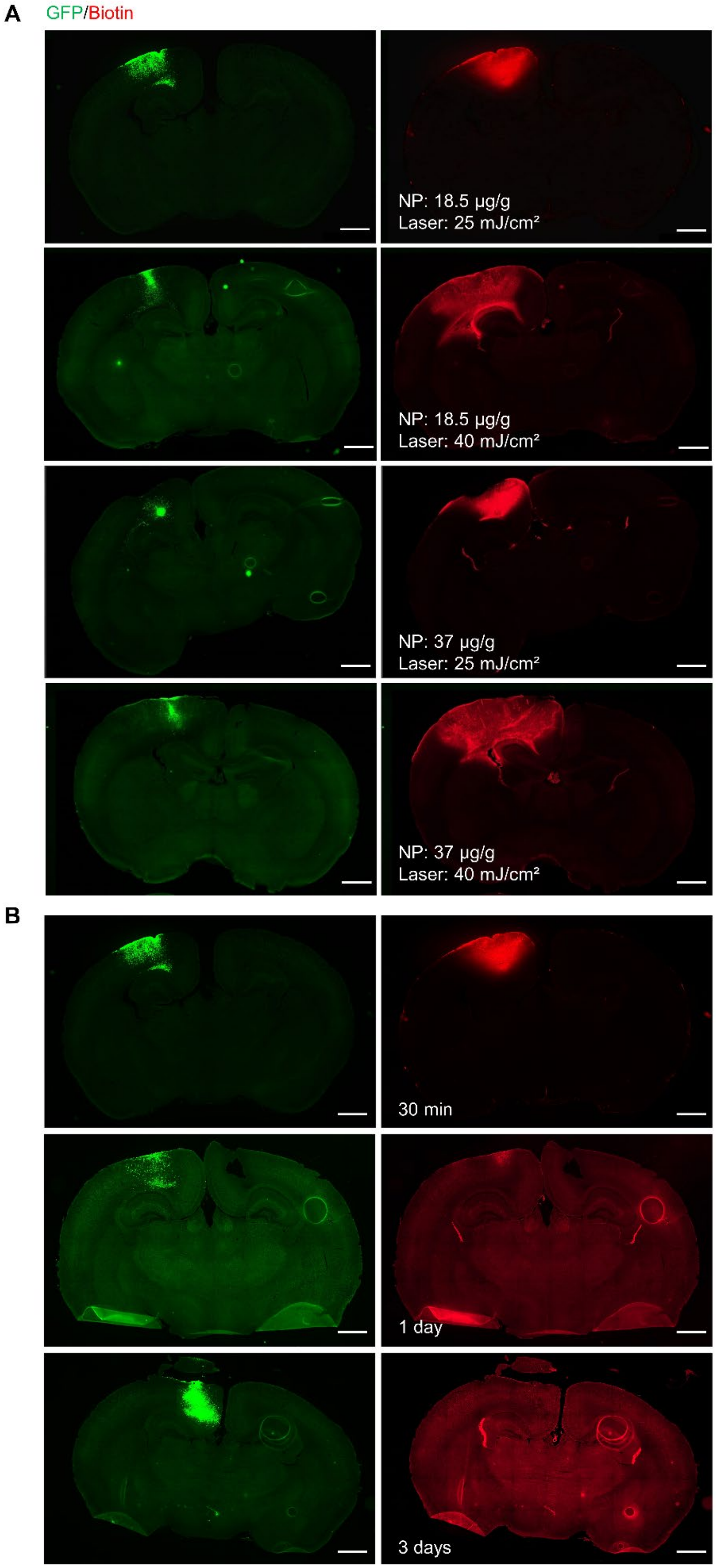
A
GFP/Biotin
NP: 18.5 µg/g
Laser: 25 mJ/cm²
NP: 18.5 µg/g
Laser: 40 mJ/cm²
NP: 37 µg/g
Laser: 25 mJ/cm²
NP: 37 µg/g
Laser: 40 mJ/cm²
B
30 min
1 day
3 days

**Fig. S3. BBTB modulation in PS5A GBM model.** (A) Optimization of optoBBTB in PS5A1 GBM model. NP injection dose was 18.5 μg/g or 37 μg/g. Laser fluence was 25 $mJ/cm^2$ or 40 $mJ/cm^2$. (B) Recovery of BBTB permeability after optoBBTB at 30 min, 1 day, and 3 days. The tumor was implanted in the left hemisphere. The green color indicates tumor cells encoded GFP, the red color indicates fluorescent EZ-link biotin. The scale bar represents 1 mm.

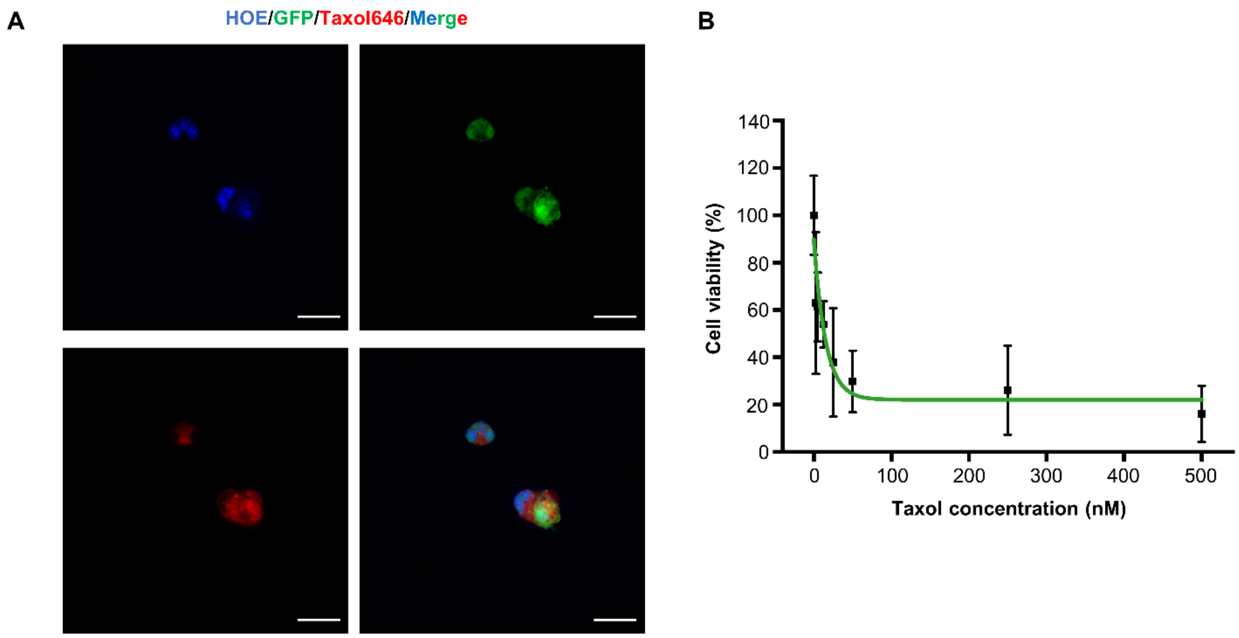


**Fig. S4. In vitro cellular uptake and cytotoxicity of fluorescent taxol in PS5A1 GBM cells.** (A) In vitro cellular uptake of fluorescent taxol. Cell nuclei, glioma cells, and fluorescent taxol are indicated in blue (HOE), green (GFP), and red colors (Taxol646). The scale bar represents 20 µm. (B) Cell viability of PS5A1 glioma cells at various concentrations (0 to 500 nM) of taxol. The IC50 value at 72 hours was 17 nM. Data were expressed as mean±SD, N=6.

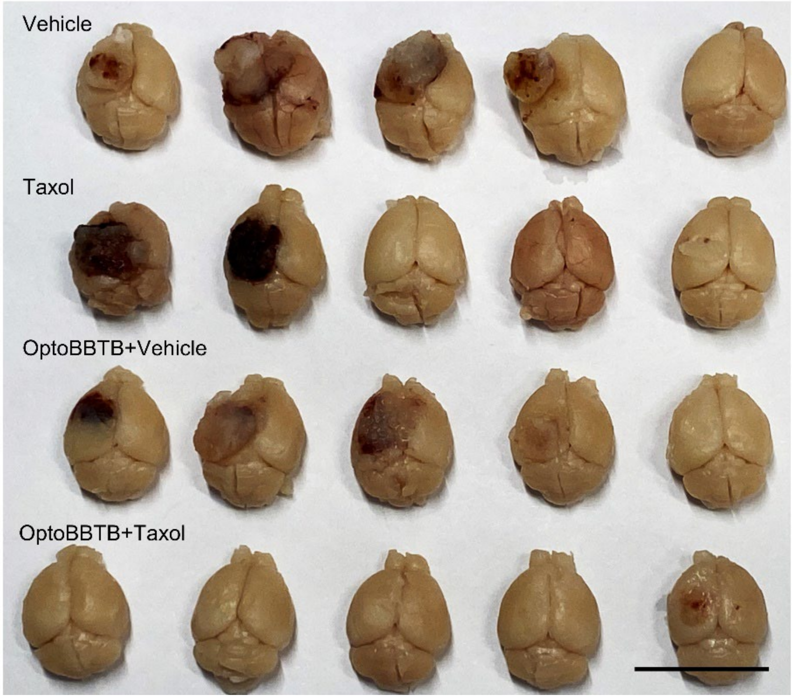


**Fig. S5. Photographs of the PS5A1 tumor size in each group at 42 dpi after 3 treatments.** The scale bar represents 1 mm.

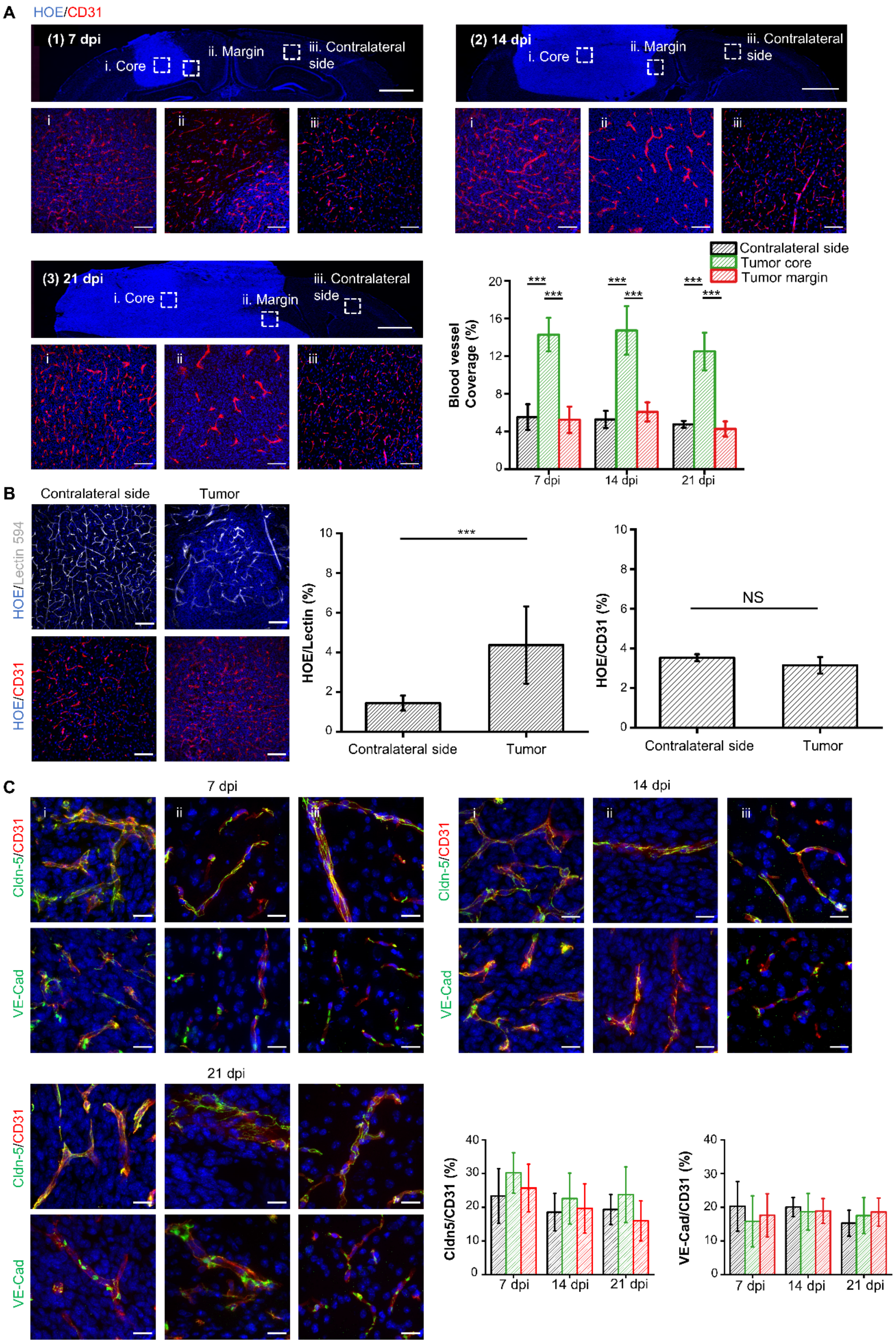
A
HOE/CD31
(1) 7 dpi
i. Core
ii. Margin
iii. Contralateral side
(2) 14 dpi
(3) 21 dpi
Contralateral side
Tumor core
Tumor margin
Blood vessel Coverage (%)
7 dpi
14 dpi
21 dpi
***
B
Contralateral side
Tumor
HOE/Lectin 594
HOE/CD31
HOE/Lectin (%)
HOE/CD31 (%)
NS
C
Cldn-5/CD31
VE-Cad
Cldn5/CD31 (%)
VE-Cad/CD31 (%)

**Fig. S6**. **Blood vessel labeling and IHC staining of junctional proteins in 73C GBM model.** (A) IHC staining and quantification of blood vessels using CD31 at 7 dpi, 14 dpi and 21 dpi. The ROIs selected are (i) tumor core, (ii) tumor margin, and (iii) contralateral side with no tumor. The scale bar represents 1 mm in the upper panel and 50 μm in the bottom panels. Quantification of blood vessel coverage was performed by CD31 area fraction. One-way Anova, ***$p<0.001$. N=15 images from 3 mice. (B) A comparison of blood vessels labeling with tomato lectin594 or CD31 at 7 dpi. The cell nuclei were labeled by Hoechst staining (HOE). Scale bar represents 100 μm. Quantification of the ratio of cell nuclei to blood vessels was performed by area fraction. Student's t-test, ***$p<0.001$. NS: no significant difference. N=10 images from one mouse. (C) IHC staining of Claudin-5 (Cldn5) and VE-Cadherin (VE-Cad) at 7 dpi, 14 dpi, and 21 dpi. The blood vessels were stained with CD31 and the cell nuclei were indicated by Hoechst staining (HOE). The ROIs selected are (i) tumor core, (ii) tumor margin, and (iii) contralateral side with no tumor. The scale bar represents 1 mm in the upper panel and 20 μm in the bottom panels. Quantification of the expression of Claudin-5 and VE-Cadherin over CD31 was analyzed by area fraction. One-way Anova, no significant difference was observed. N=15 images from 3 mice.

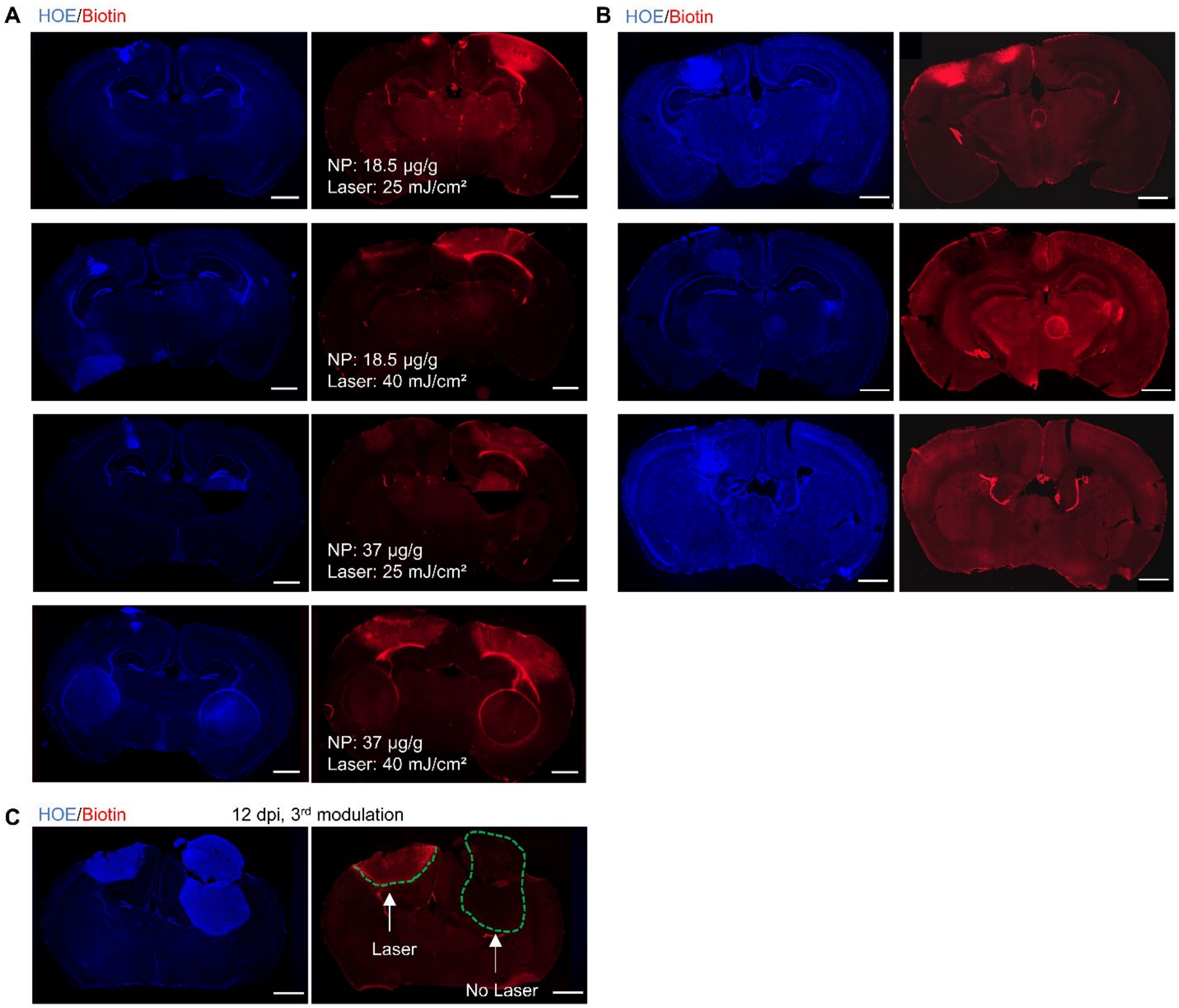


**Fig. S7. BBTB modulation in 73C GBM model.** (A) Optimization of optoBBTB in 73C GBM model. NP injection dose was 18.5 µg/g or 37 µg/g. Laser fluence was 25 mJ/cm$^2$ or 40 mJ/cm$^2$. (B) Recovery of BBTB permeability after optoBBTB at 30 min, 1 day, and 3 days. The tumor was implanted in the left hemisphere. (C) No optoBBTB modulation showed no obvious EZ-link biotin leakage into the tumor. 73C GBM cells were injected to both sides of the brain. The left hemisphere received optoBBTB. The blue color indicates Hoechst staining (HOE), the red color indicates fluorescent EZ-link biotin. The scale bar represents 1 mm.

A

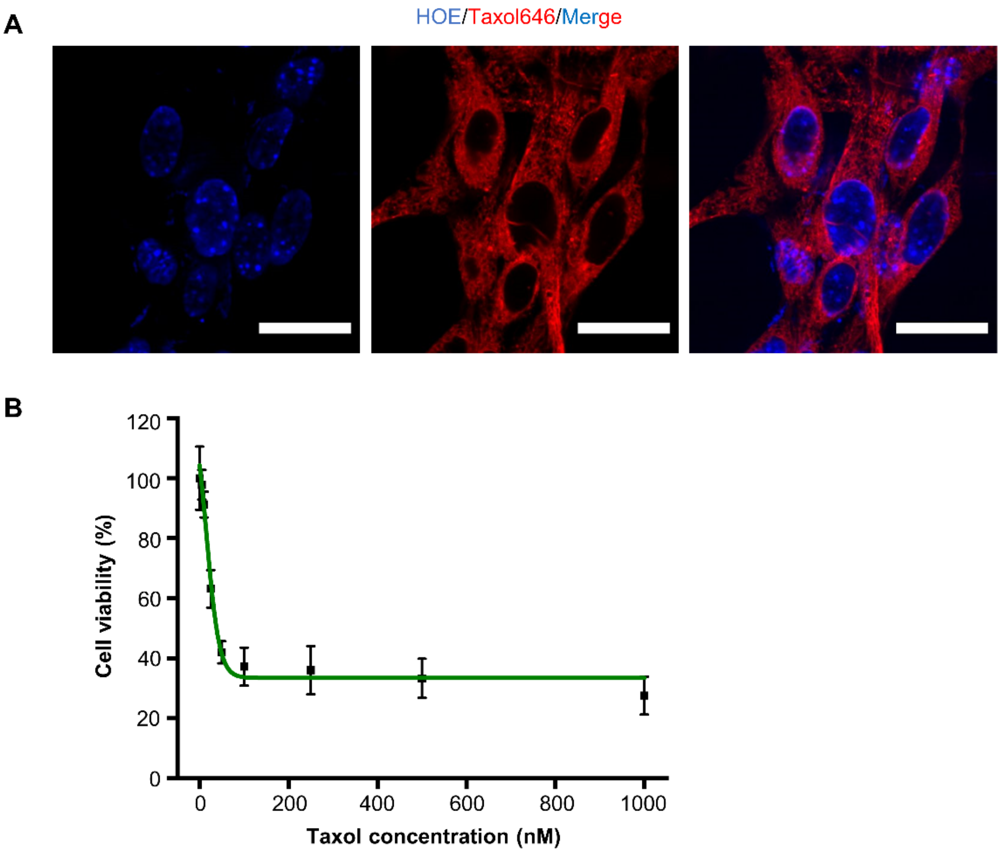


**Fig. S8. In vitro cellular uptake and cytotoxicity of fluorescent taxol646 in 73C tumor cells.** (A) In vitro cellular uptake of fluorescent taxol. Cell nuclei and fluorescent taxol are indicated in blue (HOE) and red colors (Taxol646). The scale bar represents 10 μm. (B) Cell viability of 73C tumor cells at various concentrations (0 to 1000 nM) of taxol. The IC50 value at 72 hours was 37 nM. Data were expressed as mean±SD, N=6.

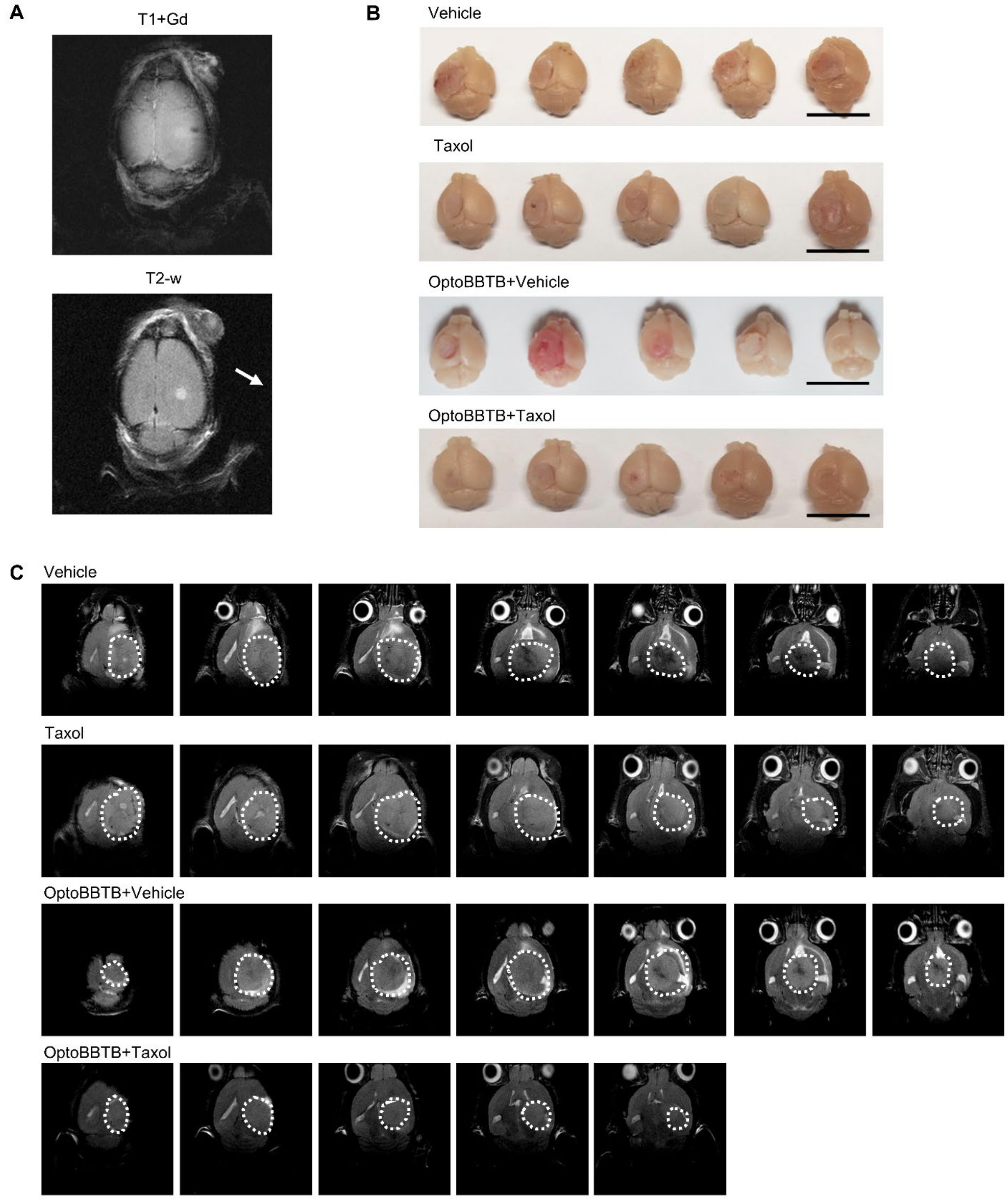


**Fig. S9. Size analysis of 73C GBM model. (A)**MRI images of 73C GBM at 3 dpi. Top, T1-weighted scan post gadolinium (Gd) injection. Bottom, T2-weighted scan. (B) Photograph of 73C tumor size at 15 dpi after 3 treatments. The scale bar represents 1 mm. (C) Representative Magnetic resonance imaging of the tumor volume in each group at 15 dpi after 3 treatments. Images were obtained from 1 mouse in each group. The tumor area was indicated by white dashed circles.

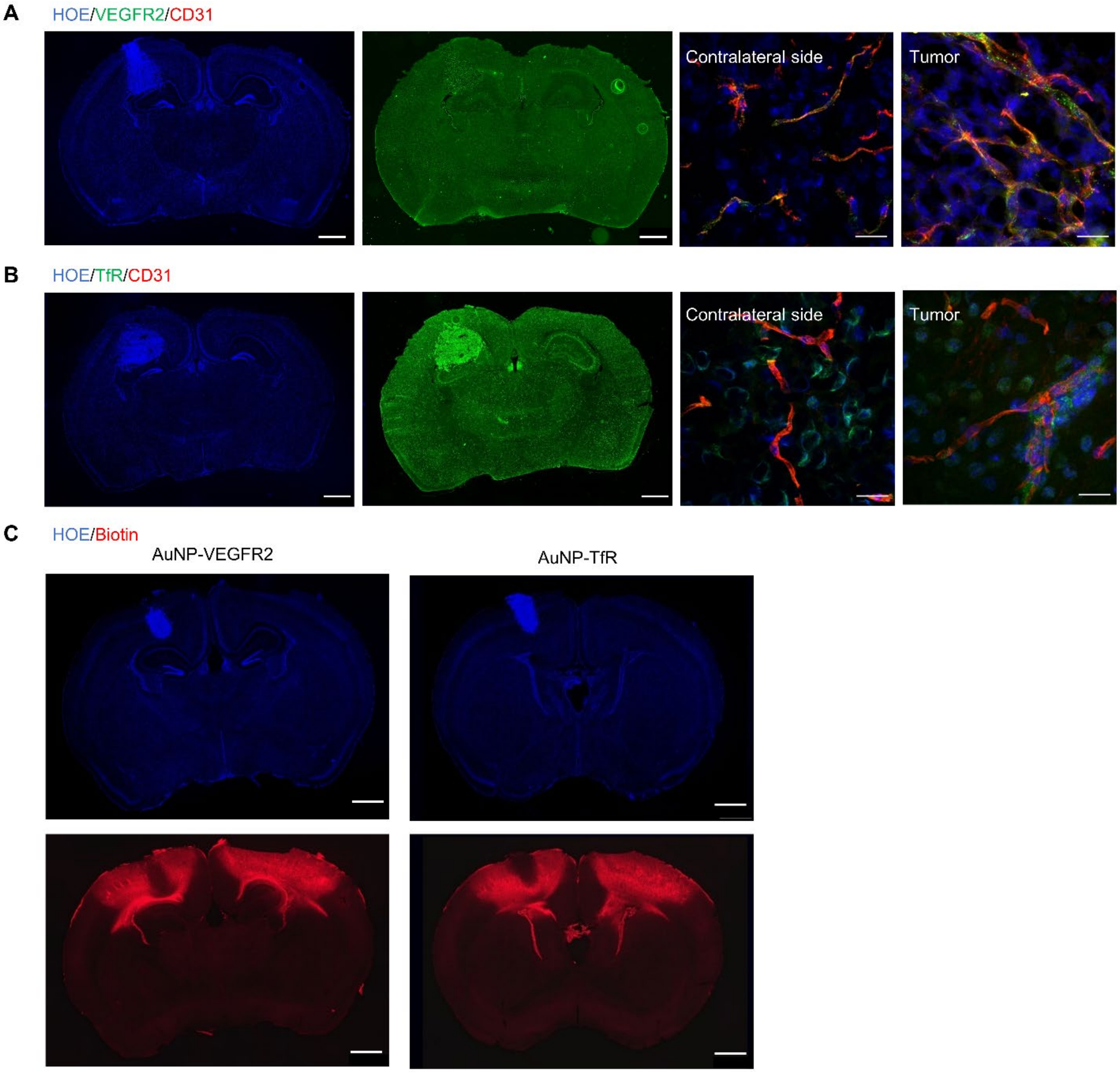


**Fig. S10. BBTB modulation using different targets in the 73C glioma mode**l. (A) IHC staining of vascular endothelial growth factor receptor 2 (VEGFR2) at 7 dpi. (B) IHC staining of transferrin receptor (TfR) at 7 dpi. The blood vessel was stained by CD31. (C) A comparison of BBTB modulation efficacy using AuNP-VEGFR2 and AuNP-TfR. The cell nuclei were indicated by Hoechst staining (HOE). The nanoparticle dose is 37 µg/g and the laser dose is 40 mJ/cm$^2$, 1 pulse. Laser was applied to both sides of the brain. The scale bar represents 1 mm in the slide scanner images and 20 µm in zoom-in images.

**Table S1**. Optimization of the optoBBTB in PS5A1 glioma-bearing mice.

| | Nanoparticle type | Nanoparticle dose | Laser dose |
|---|---|---|---|
| Condition 1 | AuNP-BV11 | 18.5 µg/g | 25 $mJ/cm^2$, 1 pulse |
| Condition 2 | AuNP-BV11 | 18.5 µg/g | 40 $mJ/cm^2$, 1 pulse |
| Condition 3 | AuNP-BV11 | 37 µg/g | 25 $mJ/cm^2$, 1 pulse |
| Condition 4 | AuNP-BV11 | 37 µg/g | 40 $mJ/cm^2$, 1 pulse |

**Table S2**. Optimization of the optoBBTB in 73C glioma-bearing mice.

| | Nanoparticle type | Nanoparticle dose | Laser dose |
|---|---|---|---|
| Condition 1 | AuNP-BV11 | 18.5 µg/g | 25 $mJ/cm^2$, 1 pulse |
| Condition 2 | AuNP-BV11 | 18.5 µg/g | 40 $mJ/cm^2$, 1 pulse |
| Condition 3 | AuNP-BV11 | 37 µg/g | 25 $mJ/cm^2$, 1 pulse |
| Condition 4 | AuNP-BV11 | 37 µg/g | 40 $mJ/cm^2$, 1 pulse |
| Condition 5 | AuNP-TfR | 37 µg/g | 40 $mJ/cm^2$, 1 pulse |
| Condition 6 | AuNP-VEGFR | 37 µg/g | 40 $mJ/cm^2$, 1 pulse |
